# Tsa1 is the dominant peroxide scavenger and a source of H_2_O_2_-dependent GSSG production in yeast

**DOI:** 10.1101/2024.07.03.601836

**Authors:** Jannik Zimmermann, Lukas Lang, Gaetano Calabrese, Hugo Laporte, Prince S Amponsah, Christoph Michalk, Tobias Sukmann, Julian Oestreicher, Anja Tursch, Esra Peker, Theresa N E Owusu, Matthias Weith, Leticia Prates Roma, Marcel Deponte, Jan Riemer, Bruce Morgan

## Abstract

Hydrogen peroxide (H_2_O_2_) is an important biological molecule, functioning both as a second messenger in cell signaling and, especially at higher concentrations, as a cause of cell damage. Cells harbor multiple enzymes that have peroxide reducing activity *in vitro*. However, the contribution of each of these enzymes towards peroxide scavenging *in vivo* is less clear. Therefore, to directly investigate *in vivo* peroxide scavenging, we used the genetically encoded peroxide sensors, roGFP2-Tsa2ΔC_R_ and HyPer7, to systematically screen the peroxide scavenging capacity of yeast thiol and heme peroxidase mutants. We show that the 2-Cys peroxiredoxin Tsa1 alone is responsible for almost all exogenous H_2_O_2_ and *tert*-butyl hydroperoxide scavenging. The two catalases and cytochrome *c* peroxidase only produce observable scavenging defects at higher H_2_O_2_ concentrations when these three heme peroxidases are deleted in combination. We also analyzed the reduction of Tsa1 *in vitro*, revealing that the enzyme is efficiently reduced by thioredoxin 1 with a rate constant of 2.8×10^6^ M^−1^s^−1^. When thioredoxins are oxidized, Tsa1 can become an important source of H_2_ O_2_ -dependent cytosolic glutathione disulfide production in yeast. Our findings clarify the importance of the various thiol and heme peroxidases for peroxide removal and suggest that most thiol peroxidases have alternative or specialized functions in specific subcellular compartments.

## Introduction

Hydrogen peroxide (H_2_O_2_) is an important cellular redox molecule. It has multiple cellular sources, including several enzymes that appear to be dedicated to the direct or indirect production of H_2_O_2_. Whilst it is now well understood that H_2_O_2_ can function as a second messenger in cellular signaling, it is also unequivocal that beyond a certain threshold, H_2_O_2_ is toxic to cells and can readily lead to cell death ^1,2^. In the context of both signaling and toxicity it is likely crucial for cells to tightly control H_2_ O_2_ levels.

Cells harbor a range of H_2_O_2_ scavenging enzymes, including thiol-or selenol-dependent peroxidases, for example, peroxiredoxins and glutathione peroxidases, as well as heme peroxidases such as catalases and cytochrome *c* peroxidases ^3-6^. Nonetheless, whilst many enzymes can efficiently reduce H_2_O_2_ *in vitro*, the contribution of a specific enzyme towards the total cellular H_2_O_2_ scavenging capacity is often less clear. It will depend upon the expression level of the according gene, the subcellular compartment, and the enzymatic activity relative to other H_2_O_2_ reactive enzymes present ^7^. Furthermore, the relevance may change dynamically over time in response to changing environmental conditions, including carbon source, culture growth phase and culture density, previous exposure to compounds including oxidants, electrophiles and heavy metals, and potentially even illumination ^8-11^. The physiological relevance and relative importance of different peroxidases also depends on the various organism-dependent functions they exert, ranging for example from peroxide detoxification and H_2_O_2_ -dependent signal transduction ^3,4,12,13^ to chaperone activity ^14,15^ or moonlighting functions such as the formation of the mitochondrial capsule of sperm cells ^16^. Despite this functional heterogeneity, the physiological relevance of peroxidases for the removal of cellular H_2_O_2_ has typically been investigated using *indirect* assays, for example, by correlating cell growth or cell death to peroxide treatments ^12,17-29^. It is frequently assumed that a difference in the sensitivity of a specific mutant strain towards H_2_O_2_, or other peroxides, is indicative of a change in the peroxide scavenging capacity of that mutant compared to wild-type cells. The validity of this approach depends upon the *assumption* that changes in H_2_O_2_ scavenging capacity always negatively correlate with sensitivity towards H_2_O_2_-induced growth arrest or cell death. However, recent studies show that peroxide scavenging capacity and sensitivity to peroxide-induced cell death can be uncoupled, indicating that the assumption is not always valid. For example, it has been reported that depletion of pools of reduced thioredoxins or glutathione (GSH) (and the accumulation of oxidized thioredoxins or glutathione disulfide) may be important mediators of H_2_O_2_ toxicity ^30-34^. In addition, misregulation of protein kinase A (PKA) signaling was proposed to underlie the sensitivity of *TSA1* knockout cells to H_2_O_2_ -induced cell death ^35^. In general, these studies question whether H_2_O_2_ scavenging activity *per se* is always inversely correlated with sensitivity towards H_2_O_2_-induced cell death or even suggest that the activity of some H_2_O_2_ removing enzymes does not prevent but rather promotes cell death, for example via depletion of reducing equivalents.

In this study we therefore sought to investigate the relative importance for peroxide removal of the various thiol and heme peroxidases in baker’s yeast, *Saccharomyces cerevisiae*. In contrast to indirect growth assays, we *directly* and *quantitatively* analyzed the peroxide scavenging capacity of deletion and overexpression mutants using two ultra-sensitive, genetically encoded H_2_O_2_ sensors, roGFP2-Tsa2ΔC_R 36_ and HyPer7 ^37^, that were localized to the mitochondrial matrix and cytosol, respectively. We found that the typical 2-Cys peroxiredoxin Tsa1 is the major scavenger of both H_2_O_2_ and *tert*-butyl hydroperoxide (*t*-BuOOH) in fermentative and respiratory conditions and efficiently outcompetes all other endogenous cellular thiols for H_2_O_2_ including the reactive thiol on the ultra-sensitive H_2_O_2_ probe, HyPer7. Regarding heme peroxidases, the two yeast catalases only have a noticeable impact on cellular H_2_O_2_ scavenging at higher H_2_O_2_ concentrations and, even then, only when the encoding genes are simultaneously deleted in combination with a deletion of the gene encoding cytochrome *c* peroxidase. Finally, we analyzed the reduction of Tsa1 *in vitro* and in yeast, revealing that the enzyme is efficiently reduced by thioredoxin 1 (Trx1) but can become a major source of H_2_O_2_-dependent glutathione disulfide (GSSG) production when thioredoxins are oxidized.

## Results

### Tsa1 is the major scavenger of cellular peroxide

We first sought to directly quantify the capacity of the yeast cytosol to eliminate H_2_O_2_ and *t*-BuOOH and to define a ‘hierarchy of importance’ of the different peroxide removing enzymes. To this end, we utilized an assay that we have previously developed in which the sensitivity of the response of a mitochondrial-localized roGFP2-Tsa2ΔC_R_ probe (Su9-roGFP2-Tsa2ΔC_R_) to exogenously applied peroxide serves as a proxy for the H_2_O_2_ scavenging capacity of the cytosol ^30,36^. The magnitude of the probe response is dependent upon the quantity of exogenous peroxide that reaches the matrix and thus is inversely dependent upon the peroxide scavenging capacity of the cytosol (Fig. 1a). Baker’s yeast harbors five peroxiredoxins, including the highly abundant Prx1-type typical 2-Cys peroxiredoxin, Tsa1, and the highly similar (86% identity) peroxiredoxin, Tsa2, as well as a Prx5-type typical 2-Cys peroxiredoxin, Ahp1, a nuclear-localized atypical 2-Cys peroxiredoxin, Dot5, and a mitochondrial matrix-localized Prx6-type 1-Cys peroxiredoxin, Prx1. Yeast also contains three glutathione peroxidase-like proteins, Gpx1, Gpx2 and Gpx3, which have active-site cysteine residues instead of selenocysteine and are predominantly reduced by thioredoxins ^38^. In addition, yeast contains two catalases, Ctt1 and Cta1, localized predominantly to the cytosol and peroxisomes respectively, as well as a mitochondrial intermembrane space-localized cytochrome *c* peroxidase, Ccp1 (Fig. 1a). We monitored Su9-roGFP2-Tsa2ΔC_R_ responses to exogenous H_2_O_2_ or *t*-BuOOH at initial concentrations ranging from 0–2 mM in wild-type cells as well as cells individually deleted for the genes encoding both catalases, cytochrome *c* peroxidase, or each of the yeast thiol peroxidases (including Gpx1, Gpx2 and Gpx3 as well as four of the five peroxiredoxins; we omitted mitochondrial Prx1 as this was analyzed recently ^30^). The response of the Su9-roGFP2-Tsa2ΔC_R_ probe was followed in a fluorescence plate-reader as described previously ^39^. For every probe response, we determined the integrated area under the background-corrected response curve between 0 and 2000 seconds (Fig. 1b). Intriguingly, our data revealed that deletion of *TSA1* led to a highly significant increase in Su9-roGFP2-Tsa2ΔC_R_ probe response to both exogenous H_2_O_2_ and *t*-BuOOH, at concentrations from 20–200 and 20–40 µM respectively, compared to wild-type cells as well as all other gene deletion mutants (P < 0.001 for all samples) (Fig. 1c–h). Associated source data and full statistical analyses are available in an online data repository accessible at (http://dx.doi.org/10.17632/62s536jngt.2). The additional deletion of the gene encoding Tsa2, to generate Δ*tsa1*Δ*tsa2* cells, made no significant difference to the Su9-roGFP2-Tsa2ΔC_R_ probe response (P = 0.999 for Δ*tsa1* vs. Δ*tsa1*Δ*tsa2*) (Fig1. c,f). At exogenous H_2_O_2_ and *t*-BuOOH concentrations above 500 µM and 100 µM respectively, no further increase in Su9-roGFP2-Tsa2ΔC_R_ response was observed for any strain as probe oxidation reached a maximum (Fig. 1c–h). Notably, at *t*-BuOOH concentrations greater than 100 µM, Su9-roGFP2-Tsa2ΔC_R_ responses decreased in all strains compared to the response observed in 100 µM *t*-BuOOH treated cells (Fig. 1f–h). This effect is consistent with hyperoxidation and therefore inactivation of the Tsa2 moiety of the Su9-roGFP2-Tsa2ΔC_R_ probe as we have shown previously ^30,36,40,41^. In summary, our data suggest that Tsa1 alone is the major cytosolic peroxide scavenger and strongly limits the diffusion of both H_2_O_2_ and *t*-BuOOH through the yeast cytosol.

**Figure 1.**
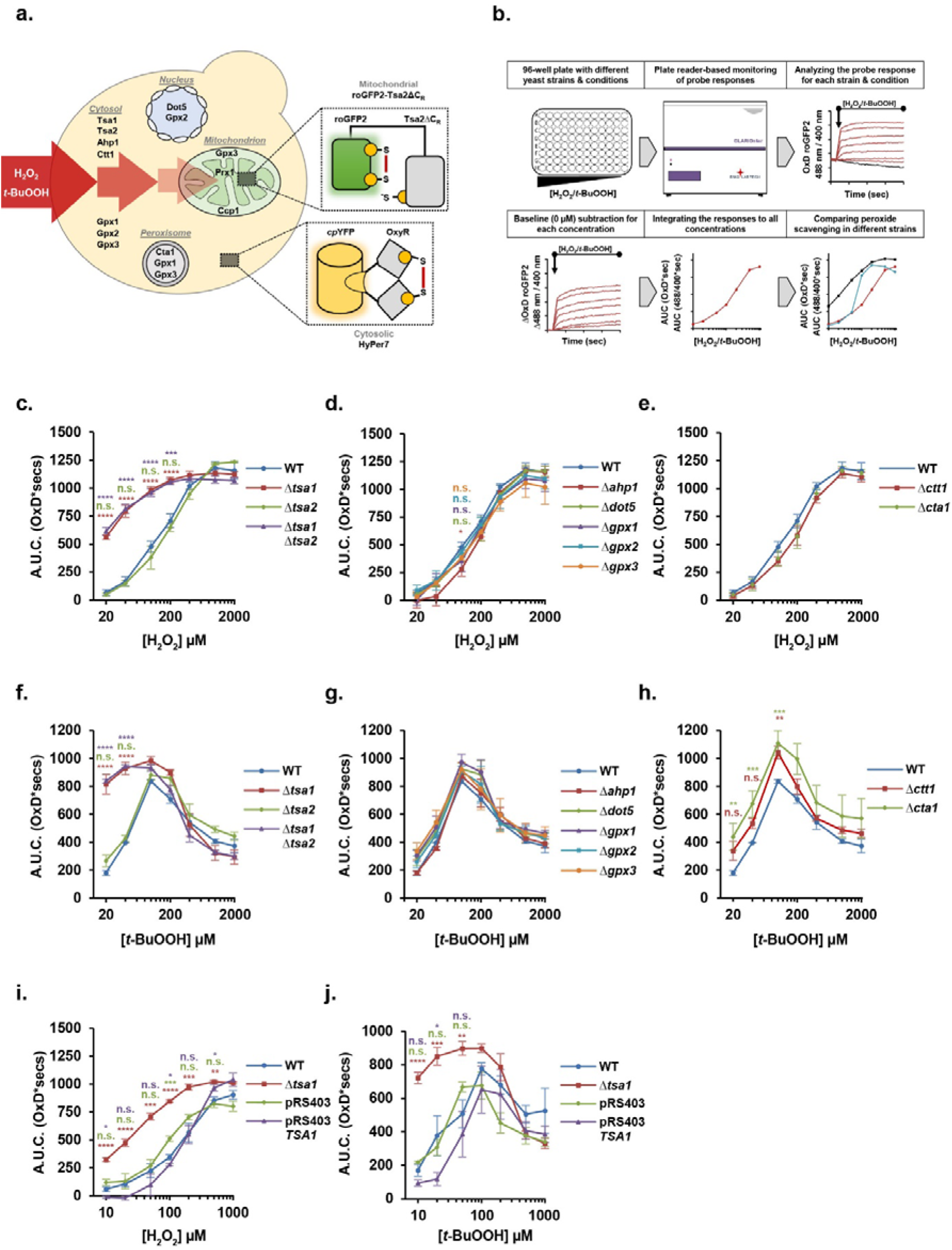
Cellular peroxide scavenging is strongly dependent upon TSA1 expression. **a**. Schematic illustration of the experimental setup. A matrix localized, ultra-sensitive, Su9-roGFP2-Tsa2ΔC_R_ peroxide sensor, was expressed in strains deleted for or overexpressing the genes encoding the various thiol peroxidases, catalases and cytochrome *c* peroxidase, either individually or in combination. The response of the Su9-roGFP2-Tsa2ΔC_R_ sensor is dependent upon the quantity of peroxide that can diffuse through the cytosol, which in turn, is inversely dependent upon the peroxide scavenging capacity. Therefore, the sensitivity of the Su9-roGFP2-Tsa2ΔC_R_ probe response is inversely dependent upon the cellular peroxide scavenging capacity. **b**. Su9-roGFP2-Tsa2ΔC_R_ probe responses were analyzed by calculating the integrated area under the curve (A.U.C.) of the untreated baseline corrected response between 0 and 2000 secs. **c,d,e**. Su9-roGFP2-Tsa2ΔC_R_ probe responses were measured in the indicated yeast deletion strains in response to exogenous H_2_O_2_ in the yeast strains indicated. The data for wild-type (WT) cells are replicated in each panel. **f,g,h**. Su9-roGFP2-Tsa2ΔC_R_ probe responses were measured in the indicated yeast deletion strains in response to exogenous *t*-BuOOH in the yeast strains indicated. The data for WT are replicated in each panel. **i,j**. Su9-roGFP2-Tsa2ΔC_R_ probe responses were measured in response to *i.* exogenous H_2_O_2_ or *j.* exogenous *t*-BuOOH applied to BY4742 wild-type or Δ*tsa1* cells, as well as BY4742 cells containing an empty genomically integrated pRS403 vector or a pRS403-*TSA1* vector, which provides cells with an additional copy of *TSA1*. Error bars in all panels represent the standard error of the mean for at least three independent repeats. P values were determined by a one-way ANOVA analysis followed by a Tukey’s (HSD) test. P values for the difference between the WT and every other strain is represented on each panel with stars. The color of the stars corresponds to the respective deletion strain being compared to WT. n.s. = not significant, * = P ≤ 0.05, ** = P ≤ 0.005, *** = P ≤ 0.001, **** = P ≤ 0.0001.

To further investigate our observations of the dominant importance of Tsa1, we next asked about the impact of overexpressing *TSA1*. To this end, we made use of a strain containing an extra genomic copy of the *TSA1* locus (generated using the integrative plasmid pRS403-*TSA1*) ^42^. We observed a trend of decreased mitochondrial Su9-roGFP2-Tsa2ΔC_R_ probe response in pRS403-*TSA1* cells, compared to wild-type and pRS403-empty controls, for both H_2_O_2_ and *t*-BuOOH at exogenous concentrations below 200 µM and 100 µM respectively (Fig. 1i,j), which was significant for some but not all concentrations of exogenous peroxide (P < 0.05). Thus, additional Tsa1 can further improve cytosolic peroxide removal.

Although Tsa1 was originally identified and purified as the dominant scavenger of peroxides in yeast ^43^ in accordance with our non-invasive intracellular measurements, it was surprising to us that we detected almost no effect of deleting the genes encoding any other thiol peroxidase or catalase. We thus asked if compensatory upregulation, perhaps including upregulation of Tsa1, may mask the effect of deleting other thiol peroxidases. To address this possibility, we investigated whether additional deletion of the peroxiredoxin genes *TSA2, AHP1* or *DOT5* in a Δ*tsa1* background would further impair H_2_O_2_ or *t*-BuOOH scavenging. Comparing Δ*tsa1* cells with Δ*tsa1*Δ*tsa2*, Δ*tsa1*Δ*ahp1*, Δ*tsa1*Δ*dot5*, Δ*tsa1*Δ*tsa2*Δ*ahp1* and Δ*tsa1*Δ*tsa2*Δ*dot5* cells, we observed no significant difference in the Su9-roGFP2-Tsa2ΔC_R_ probe response at any concentration of H_2_O_2_ or *t*-BuOOH tested (Fig. 2a,b). For *t*-BuOOH-treated cells, in the Δ*tsa1* background, we observed an almost maximal mitochondrial Su9-roGFP2-Tsa2ΔC_R_ probe response even at the lowest *t*-BuOOH concentration tested, i.e. 10 µM. Therefore, we repeated the experiment with exogenous *t*-BuOOH concentrations from 0.1–10 µM (Fig. S1a). This experiment also revealed no significant difference between Δ*tsa1* cells and Δ*tsa1*Δ*tsa2*, Δ*tsa1*Δ*ahp1*, Δ*tsa1*Δ*dot5*, Δ*tsa1*Δ*tsa2*Δ*ahp1* or Δ*tsa1*Δ*tsa2*Δ*dot5* cells at any *t*-BuOOH concentration tested, except for Δ*tsa1* vs. Δ*tsa1*Δ*ahp1* at 10 µM (P = 0.03). In summary, deletion of *TSA1* alone significantly impairs cytosolic peroxide scavenging and further removal of other cytosolic peroxiredoxins does not exacerbate this phenotype.

**Figure 2.**
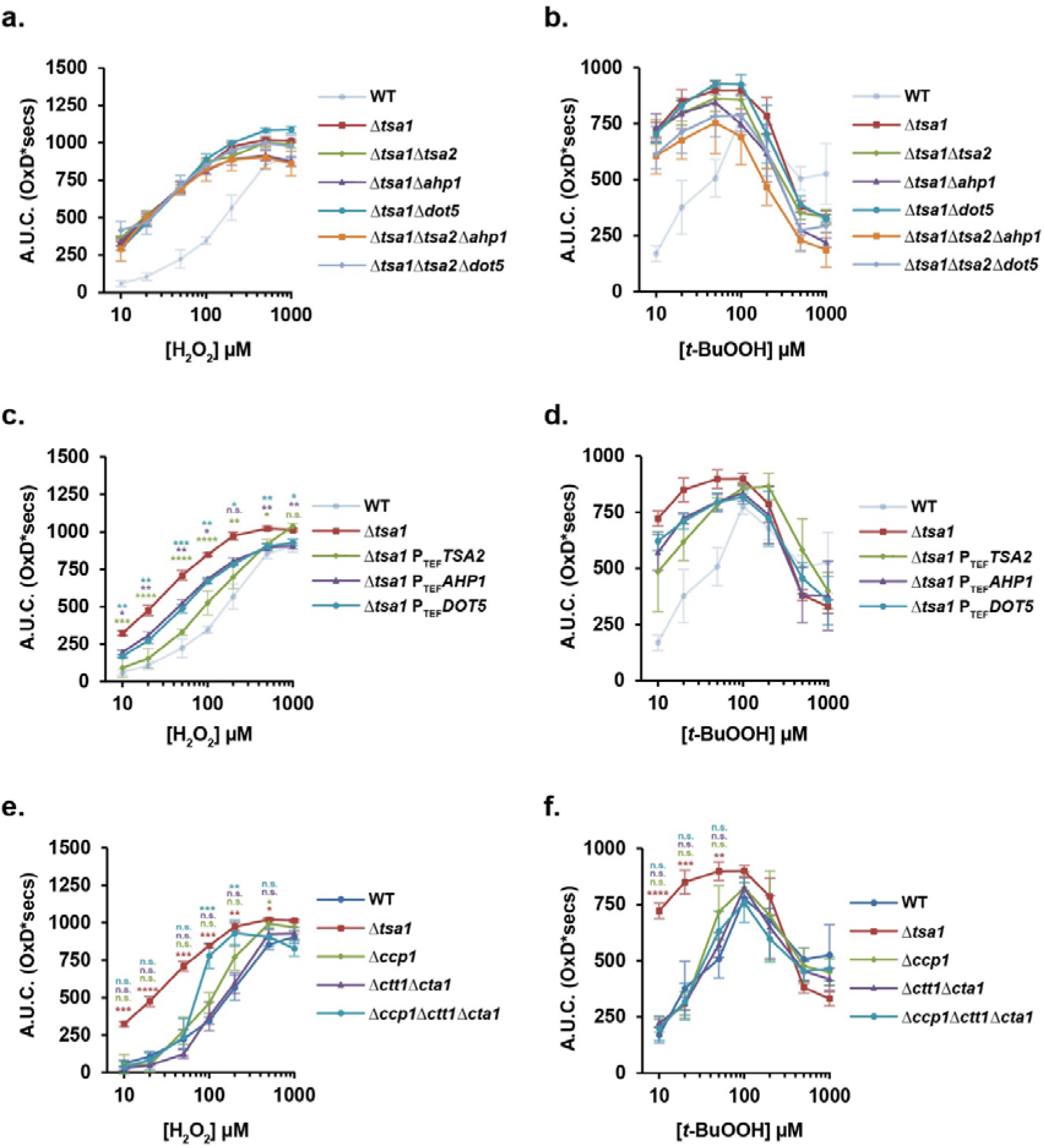
Additional deletion of TSA2, AHP1 or DOT5 in a Δtsa1 background does not further decrease cellular peroxide scavenging. **a–f**. Su9-roGFP2-Tsa2ΔC_R_ probe responses were measured in the indicated yeast deletion and overexpression strains, in response to **a,c,e.** exogenous H_2_O_2_ or **b,d,f.** exogenous *t*-BuOOH applied at the indicated concentrations. The WT and Δ*tsa1* data for H_2_O_2_ treatment are replicated in **Fig. 1i**, and **Fig. 2a,c,e.** The WT and Δ*tsa1* data for *t*-BuOOH treatment are replicated in **Fig. 1j**, and **Fig. 2b,d,f.** Error bars in all panels represent the standard error of the mean for at least three independent repeats. P values were determined by a one-way ANOVA analysis followed by a Tukey’s (HSD) test. P values for the difference between the Δ*tsa1* and every other strain (**a–d**) or between WT and every other strain (**e,f**) is represented on each panel with stars. The color of the stars corresponds to the respective deletion strain being compared to WT. n.s. = not significant, * = P ≤ 0.05, ** = P ≤ 0.005, *** = P ≤ 0.001, **** = P ≤ 0.0001.

Finally, we asked whether the overexpression of the genes encoding other cytosolic peroxiredoxins could improve cytosolic peroxide scavenging in a Δ*tsa1* background. To this end, we replaced the endogenous promoters of *TSA2, AHP1* or *DOT5* with a strong, constitutive TEF promoter in their genomic loci ^44^ to generate Δ*tsa1*P_TEF_*TSA2*, Δ*tsa1*P_TEF_*AHP1* and Δ*tsa1*P_TEF_*DOT5* strains. We again monitored mitochondrial Su9-roGFP2-Tsa2ΔC_R_ probe responses to exogenous H_2_O_2_ or *t*-BuOOH at concentrations from 0–1 mM. Interestingly, we observed a significant decrease in Su9-roGFP2-Tsa2ΔC_R_ probe response in Δ*tsa1* P_TEF_*TSA2*, Δ*tsa1* P_TEF_*AHP1* and Δ*tsa1* P_TEF_*DOT5* cells compared to Δ*tsa1* cells at all H_2_O_2_ concentrations from 0–200 µM (Fig. 2c). In response to *t*-BuOOH we observed a trend towards decreased Su9-roGFP2-Tsa2ΔC_R_ responses in the Δ*tsa1*P_TEF_*TSA2*, Δ*tsa1*P_TEF_*AHP1* and Δ*tsa1*P_TEF_*DOT5* strains compared to Δ*tsa1* at low *t-*BuOOH concentrations, however, these differences were not statistically significant (P > 0.05, Fig. 2d). At lower *t*-BuOOH concentrations, from 0.1–20 µM, we again observed a trend of decreased Su9-roGFP2-Tsa2ΔC_R_ probe response in Δ*tsa1* P_TEF_*TSA2*, Δ*tsa1*P_TEF_*AHP1* and Δ*tsa1*P_TEF_*DOT5* cells compared to Δ*tsa1* cells, but this was not significant at any concentration tested (P > 0.05, Fig. S1b). In summary, overexpression of *TSA2, AHP1* or *DOT5* increases cytosolic H_2_O_2_ scavenging in Δ*tsa1* cells but does not significantly increase *t*-BuOOH scavenging.

### Cytochrome c peroxidase and catalases are collectively important at high H_2_O_2_ concentrations

In contrast to thiol peroxidases, which usually have micromolar *K*_m_^*app*^ values, catalases use H_2_O_2_ as an oxidant *and* as a reductant, show no real substrate saturation, and do not require NADPH or a thiol adapter as electron donor during the regular catalytic cycle ^45,46^. Therefore, it is thought that catalases are more important for scavenging at higher concentrations of H_2_O_2_, e.g., when thiol peroxidases are apparently saturated or inactivated by hyperoxidation, or when the NADPH-dependent replenishment of thiol pools might become rate-limiting ^7,47^. Nonetheless, as shown in Fig. 1, individual deletion of either *CTA1* or *CTT1*, had no significant impact on scavenging at any H_2_O_2_ concentration tested. Importantly, catalases are unable to catalyze *t*-BuOOH reduction, which thus serves as a negative control for further experiments. We therefore asked whether deletion of *CTA1* and *CTT1* in combination would have an impact on cytosolic H_2_O_2_ scavenging. Furthermore, we investigated the relevance of the mitochondrial intermembrane space-localized cytochrome *c* peroxidase, Ccp1. We observed no difference in mitochondrial Su9-roGFP2-Tsa2ΔC_R_ probe response between wild-type cells and Δ*ccp1*, Δ*ctt1*Δ*cta1*, or Δ*ccp1*Δ*ctt1*Δ*cta1* cells at H_2_O_2_ concentrations from 10–50 µM (Fig. 2e). However, at higher H_2_O_2_ concentrations, i.e. 100 and 200 µM, we observed a significant increase in Su9-roGFP2-Tsa2ΔC_R_ probe response in Δ*ccp1*Δ*ctt1*Δ*cta1* cells compared to wild-type cells (P < 0.005). The Su9-roGFP2-Tsa2ΔC_R_ probe response at this concentration was similarly increased as in Δ*tsa1* cells. These observations thus suggest that Ccp1, Cta1 and Ctt1 in combination have a similar impact on scavenging of higher concentrations of H_2_O_2_ as Tsa1 but that, even in combination, they are not relevant at lower H_2_O_2_ concentrations. In contrast, we observed no significant difference in the response of the Su9-roGFP2-Tsa2ΔC_R_ probe to *t*-BuOOH in any strain compared to wild-type cells, except for the Δ*tsa1* control (Fig. 2f). Thus, our results indicate that catalases and cytochrome *c* peroxidase are only important in combination for removal of higher concentrations of H_2_O_2_, which seems consistent with the kinetic parameters and substrate specificity of catalases and cytochrome *c* peroxidase.

### HyPer7 responds specifically to H_2_O_2_ but poorly to t-BuOOH

Thus far, all our measurements were performed in cells grown with a non-fermentable glycerol/ethanol mix as the carbon source. However, yeast cells grown on high glucose are known to repress mitochondrial biogenesis as well as repress the general stress response pathway ^48-50^. We were thus interested to see if our observations were reproducible in cells grown on glucose as the sole carbon source, particularly when cells were harvested at a low cell density, i.e. when the glucose concentration in the medium is still high. Our Su9-roGFP2-Tsa2ΔC_R_-based assay is not compatible with cells grown in glucose as, due to repression of mitochondrial biogenesis, the matrix-localized Su9-roGFP2-Tsa2ΔC_R_ fluorescence is too low for reliable measurements. We therefore turned to an alternative assay, namely expressing genetically encoded H_2_O_2_ sensors in the cytosol to study the ability of endogenous H_2_O_2_ scavenging enzymes to compete with the probe for H_2_O_2_. We chose not to use roGFP2-Tsa2ΔC_R_ for these studies as we have previously shown that this probe forms heterooligomers with endogenous Tsa1 ^36^. We wanted to avoid any possibility for our results to be confounded by possible differences in probe behavior dependent upon whether it could co-assemble with endogenous Tsa1 or not. Instead, we turned to the ultrasensitive H_2_O_2_ probe HyPer7 ^37^. Preliminary characterization revealed that HyPer7, expressed in wild-type cells, responds less sensitively to exogenous H_2_O_2_ than roGFP2-Tsa2ΔC_R_ (Fig. S2a,c). Whilst cytosolic roGFP2-Tsa2ΔC_R_ responds to as little as 10 µM exogenous H_2_O_2_, HyPer7 showed no response until exogenous H_2_O_2_ was applied at a concentration of 50 µM or higher. Interestingly, we observed that HyPer7 was barely responsive to *t*-BuOOH, which is consistent with previous *in vitro* data ^37^, whereas roGFP2-Tsa2ΔC_R_ responded to *t*-BuOOH at a concentration of 10 µM or above (Fig. S2b,d). Thus, for all further experiments we limited ourselves to monitoring the response to H_2_O_2_ alone. HyPer7 containing a mutation of the reactive cysteine, HyPer7 C121S, was completely unresponsive to either H_2_O_2_ or *t*-BuOOH at any concentration tested and served as a negative control (Fig. S2e,f). Furthermore, we investigated the relative importance of the cytosolic thioredoxin and glutaredoxin systems for HyPer7 reduction. We observed that both the steady-state HyPer7 fluorescence excitation ratio, as well as the response to exogenous H_2_O_2_ was strongly increased in Δ*trx1*Δ*trx2* cells in comparison to wild-type cells, whilst we observed no difference between wild-type and Δ*grx1*Δ*grx2* cells (Fig. S3a–d). We thus conclude that HyPer7 is predominantly reduced by cytosolic thioredoxins, confirming previous reports ^51,52^, and in contrast to the GSH/glutaredoxin-dependent reduction of roGFP2-Tsa2ΔC_R 36_.

### Tsa1 scavenges the majority of H_2_O_2_ in the cytosol of cells grown with glucose as carbon source

Satisfied that HyPer7 specifically responds to H_2_O_2_ in our yeast system, we next turned to using it to address the relative importance of the various H_2_O_2_ scavengers in glucose-grown cells. Consistent with our Su9-roGFP2-Tsa2ΔC_R_ data, we show that cytosolic HyPer7 is much more efficiently oxidized by H_2_O_2_ in Δ*tsa1* cells as compared to wild-type cells (Fig. 3a). The HyPer7 response was not significantly increased in any other thiol peroxidase deletion mutant when compared to wild-type cells at any H_2_O_2_ concentration tested (Fig. 3a,b). Intriguingly, in the Δ*tsa1* cells, the HyPer7 response to H_2_O_2_ is comparable to that of roGFP2-Tsa2ΔC_R_, with HyPer7 responding to as little as 10 µM exogenous H_2_O_2_ (compare Fig. 1c and Fig. 3a). We interpret this to mean that endogenous Tsa1, due to its high concentration and high H_2_O_2_ reactivity, efficiently outcompetes HyPer7 for H_2_O_2_. In the absence of Tsa1, HyPer7 becomes the most abundant and efficient H_2_O_2_ scavenger in the cell and can then effectively compete for H_2_O_2_. Finally, we monitored the HyPer7 response to H_2_O_2_ in Δ*ccp1*, Δ*cta1*, Δ*ctt1* and Δ*ccp1*Δ*ctt1*Δ*cta1* cells in comparison to wild-type cells. Consistent with our Su9-roGFP2-Tsa2ΔC_R_ results, we observed no significant impact of catalase or cytochrome *c* peroxidase deletion on HyPer7 response at lower concentrations of exogenous H_2_O_2_ (Fig. 3c). However, we did observe a significant increase in HyPer7 response in Δ*ccp1*Δ*ctt1*Δ*cta1* cells in comparison to wild-type cells at 500 µM exogenous H_2_O_2_ (P = 0.0005). An additional *TSA1* locus decreased the HyPer7 response comparable to the roGFP2-Tsa2ΔC_R_ response in Fig. 1i (Fig. 3d). Similar effects as observed for all deletion strains in Fig. 3a–d were also seen in cells grown on non-fermentable glycerol/ethanol medium (Fig. S4). In summary, our observations of HyPer7 responses in the cytosol of glucose and glycerol/ethanol-grown cells support the conclusion that Tsa1 is the dominant scavenger of H_2_O_2_ under both fermentative and respiratory conditions. Catalases and cytochrome *c* peroxidase only have a significant impact on cellular H_2_O_2_ scavenging capacity when deleted in combination and even then, are only important at higher H_2_O_2_ concentrations.

**Figure 3.**
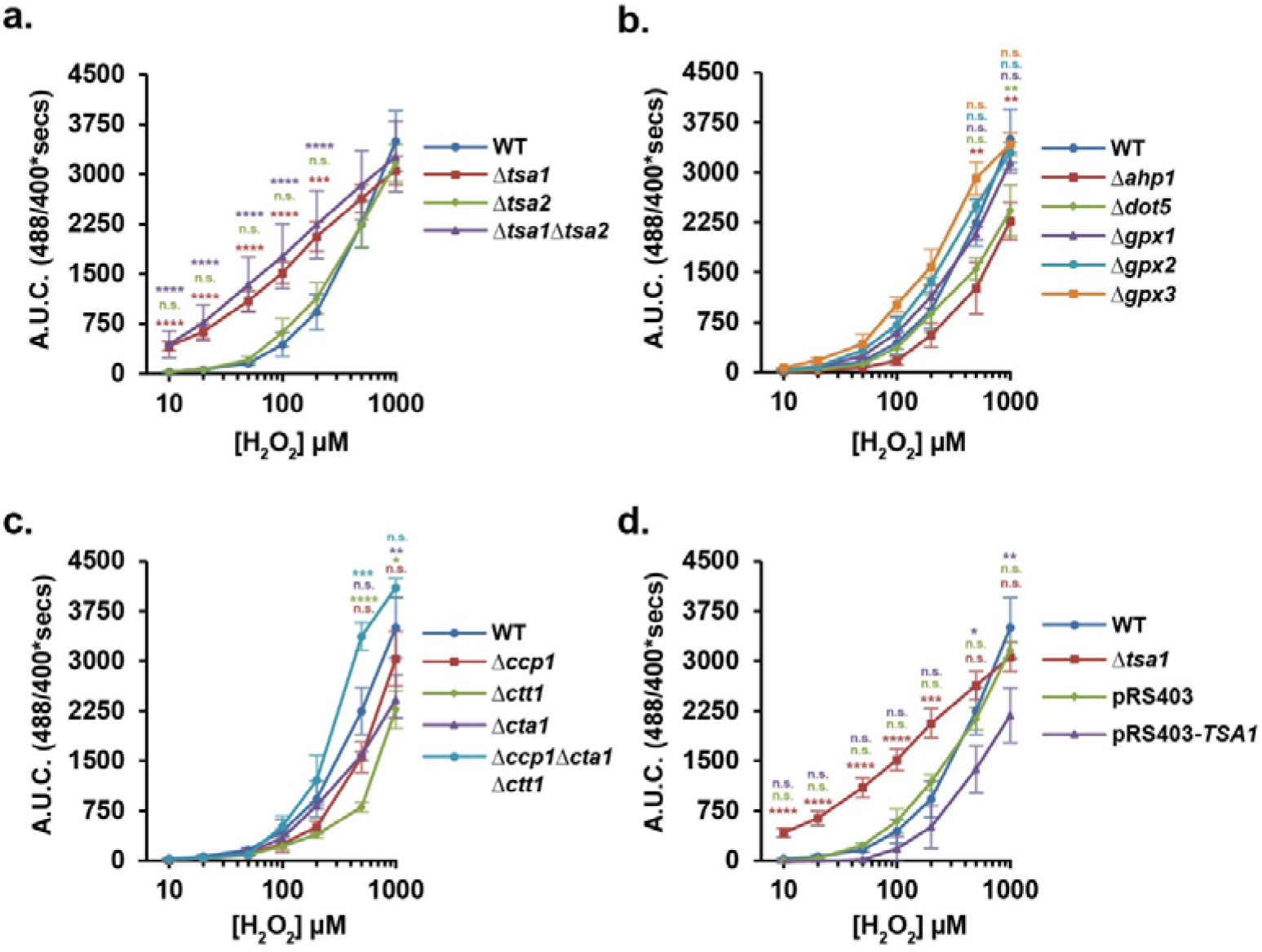
Cytosolic HyPer7 enables measurements of the cellular peroxide scavenging capacity in glucose containing media. **a–d**. HyPer7 responses were measured in the indicated yeast deletion strains in response to exogenous H_2_O_2_ applied at the indicated concentrations. The integrated A.U.C. was calculated for the first 2000 secs of the probe response. The data for wild-type (WT) cells are replicated in each panel. The data for Δ*tsa1* cells is replicated in panels **a**. and **d**. Error bars in both panels represent the standard error of the mean for at least three independent repeats. P values were determined by a one-way ANOVA analysis followed by a Tukey’s (HSD) test. P values for the difference between the WT and every other strain is represented on each panel with stars. The color of the stars corresponds to the respective deletion strain being compared to WT. n.s. = not significant, * = P ≤ 0.05, ** = P ≤ 0.005, *** = P ≤ 0.001, **** = P ≤ 0.0001.

### TSA1 deletion induces compensatory upregulation of pentose phosphate pathway enzymes

We next sought to understand whether TSA1 deletion resulted in proteome rewiring. To this end, we performed label-free quantitative proteome analysis of wild-type and Δ*tsa1*Δ*tsa2* cells. We observed large proteomic changes, with 193 and 145 proteins being significantly more or less abundant respectively, in the Δ*tsa1*Δ*tsa2* strain (Fig. S5). GO term overrepresentation analysis for up-regulated proteins indicated changes in carbohydrate metabolism in response to reactive oxygen species. In particular, glucose-6-phosphate metabolism, which is required to fuel the pentose phosphate pathway (PPP) – the most important source of cytosolic NADPH — was among the most significant terms and several PPP enzymes were among the most strongly up-regulated proteins (Fig. S5). No significantly enriched terms were detected for down-regulated proteins.

### Tsa1 is a major source of cellular glutathione peroxidase activity

The addition of H_2_O_2_ to budding yeast leads to the production of cellular GSSG, which is particularly evident in cells deleted for *GLR1*, which encodes glutathione reductase ^53^. However, the enzyme(s) responsible for the glutathione peroxidase-like activity in yeast remain unclear. Yeast has three proteins, Gpx1, Gpx2 and Gpx3, that are structurally related to mammalian glutathione peroxidases ^20,22,24^. Although glutathione-dependent activities were reported for the yeast glutathione peroxidase isoforms ^20,22,24^, no rate constants were determined, and Gpx3 was shown to be much more active with reduced thioredoxins than GSH ^38^, which is in accordance with GSH-dependent mammalian glutathione peroxidases being rather the exception ^54^. We recently demonstrated that the peroxiredoxin Prx1 is an important source of GSSG in the mitochondrial matrix upon H_2_O_2_ treatment although the impact upon total cellular GSSG levels is minor ^30^. Given the dominant role of Tsa1 in H_2_O_2_ reduction, coupled with the recent observation that GSH can serve as an electron donor for the reduction of human PRDX2 ^55^, we asked whether Tsa1 is involved in H_2_ O_2_ -dependent GSSG production.

To address this question, we used a roGFP2-Grx1 probe ^56^ to monitor the response of the cytosolic glutathione steady-state redox potential to exogenous H_2_O_2_ addition. We performed these experiments in cells deleted for *GLR1*, the gene encoding glutathione reductase, to sensitize the glutathione pool to oxidation. In this background we additionally deleted *TSA1* and/or *TSA2* to generate Δ*glr1*, Δ*glr1*Δ*tsa1, glr1*Δ*tsa2* and Δ*glr1*Δ*tsa1*Δ*tsa2* cells. Intriguingly, we observed a strong and significantly decreased probe response in both Δ*glr1*Δ*tsa1* and Δ*glr1*Δ*tsa1*Δ*tsa2* cells compared to Δ*glr1*, indicating less GSSG formation, despite the much less efficient H_2_O_2_ scavenging in the absence of Tsa1 (Fig. 4). We observed no significant difference in glutathione probe response between Δ*glr1* and Δ*glr1*Δ*tsa2* cells (Fig. 4). To exclude that the results are a *GLR1* deletion-dependent phenotype, we also monitored the responses of Grx1-roGFP2 ^56^ and roGFP2-Tsa2ΔC_R_ probes in wild-type and Δ*tsa1*Δ*tsa2* cells to monitor the response of the cytosolic glutathione steady-state redox potential and H_2_O_2_ level to exogenous H_2_O_2_ addition (Fig. S6). Consistent with the data in Fig. 4, we saw a strongly decreased Grx1-roGFP2 response and an increased roGFP2-Tsa2ΔC_R_ response in Δ*tsa1*Δ*tsa2* cells compared to wild-type cells. Our results indicate that Tsa1 can use GSH as either a direct or indirect reductant. Thus, under the chosen assay conditions, Tsa1 is an important source of H_2_O_2_-dependent GSSG production.

**Figure 4.**
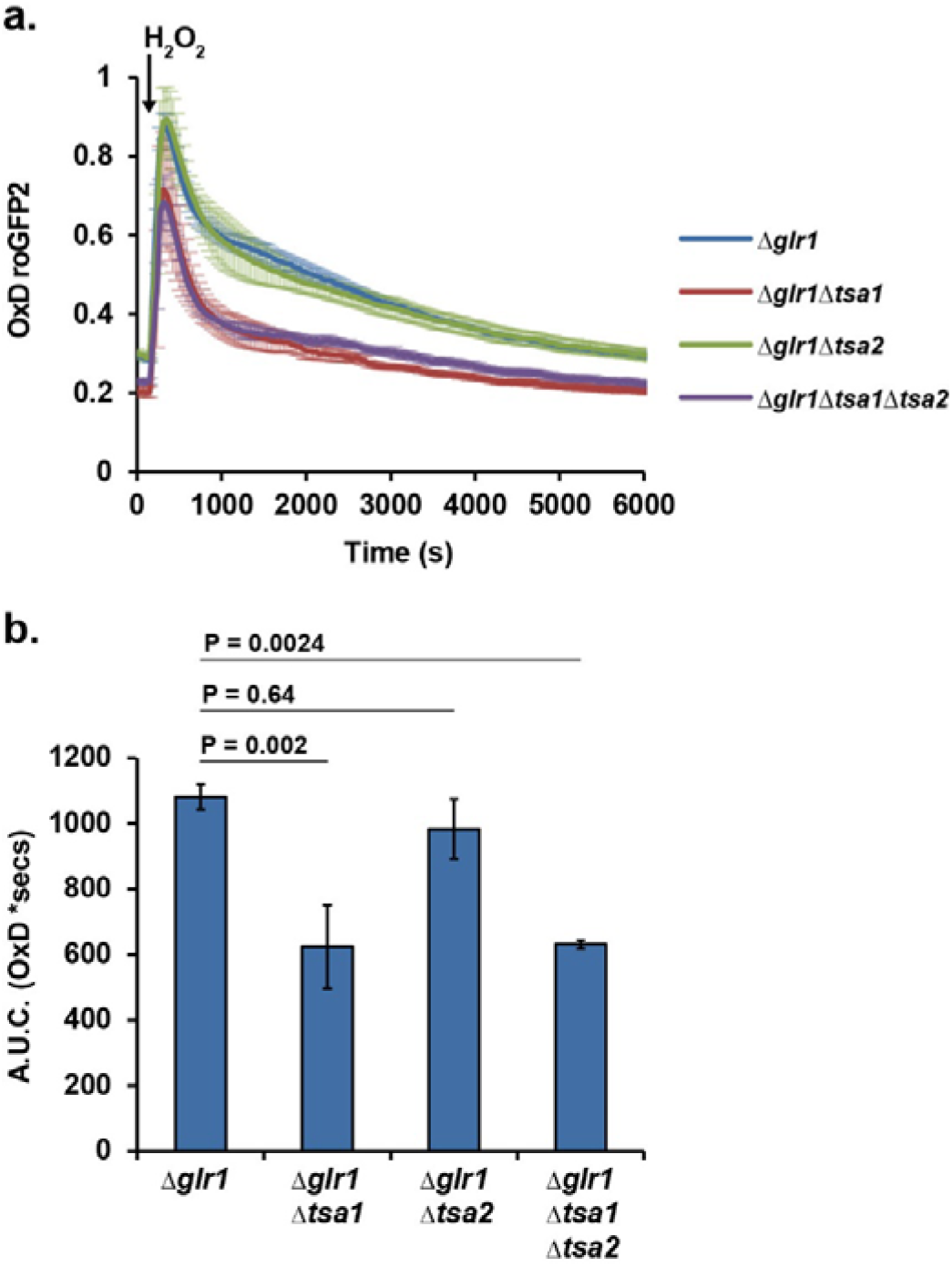
Tsa1 is a major source for cytosolic GSSG. **a.** Cytosolic roGFP2-Grx1 responses measured in BY4742 wild-type, Δ*glr1*, Δ*glr1*Δ*tsa1*, Δ*glr1*Δ*tsa2* and Δ*glr1*Δ*tsa1*Δ*tsa2* cells in response to the addition of H_2_O_2_ at the indicated time-point. **b.** A.U.C. of the roGFP2-Grx1 responses shown in panel *a.* for the first 6000 secs. Error bars in all panels represent the standard error of the mean for at least three independent repeats. P values were determined by a one-way ANOVA analysis followed by a Tukey’s (HSD) test.

### GSH slowly reduces Tsa1 in vitro

To assess whether cytosolic thioredoxins, glutaredoxins and/or GSH are efficient direct reductants of Tsa1, we determined the according rate constants in stopped-flow kinetic measurements (Figs. 5–7). To confirm our experimental setup, we mixed reduced Tsa1 in the first syringe with H_2_O_2_ in the second syringe revealing two or three distinct kinetic phases based on the altered tryptophan fluorescence of the enzyme as reported previously for Prx1-type typical 2 Cys peroxiredoxins ^57-60^ (Fig. 5a,b). A rapid decrease in fluorescence during the first phase depended on the H_2_O_2_ concentration and usually occurred during the dead-time of the measurement at higher H_2_O_2_ concentrations (Fig. 5b). This reaction phase reflects the sulfenic acid formation of the peroxidatic cysteinyl residue (Fig. 5a). The increase in fluorescence during the second phase and decrease in fluorescence during the third phase did not depend on the H_2_O_2_ concentration (Fig. 5b). These phases can be assigned to local protein unfolding and intersubunit disulfide bond formation between the peroxidatic and resolving cysteinyl residue ^58-60^. A slow fourth and a very slow fifth phase were only detected when GSH was added to reduced Tsa1 in the first syringe (Fig. 5c,d). The observed rate constant (*k*_*obs*_) for the increase in fluorescence during the fourth phase depended on the GSH concentration. A rather small second order rate constant *k*_GSH_ of 2.9 M^−1^s^−1^ and a GSH-independent rate constant of 0.010 s^−1^ were determined from the slope and y-axis intercept of a secondary plot (Fig. 5e). We assigned *k*_*GSH*_ to the nonenzymatic reduction of the intersubunit disulfide bond by GSH, whereas the GSH-independent rate constant might reflect a conformational change, the reverse reaction or a transfer of the glutathione moiety between the peroxidatic and resolving cysteinyl residue. The strong increase in fluorescence towards the potential initial fluorescence of the experiment suggests that the local refolding of Tsa1 (which necessitates a reduced and not a glutathionylated peroxidatic cysteinyl residue) also occurs during the fourth phase (Fig. 5c,d). However, this step is kinetically masked by the rate-limiting nonenzymatic reduction of the intersubunit disulfide bond (Fig. 5a).

**Figure 5.**
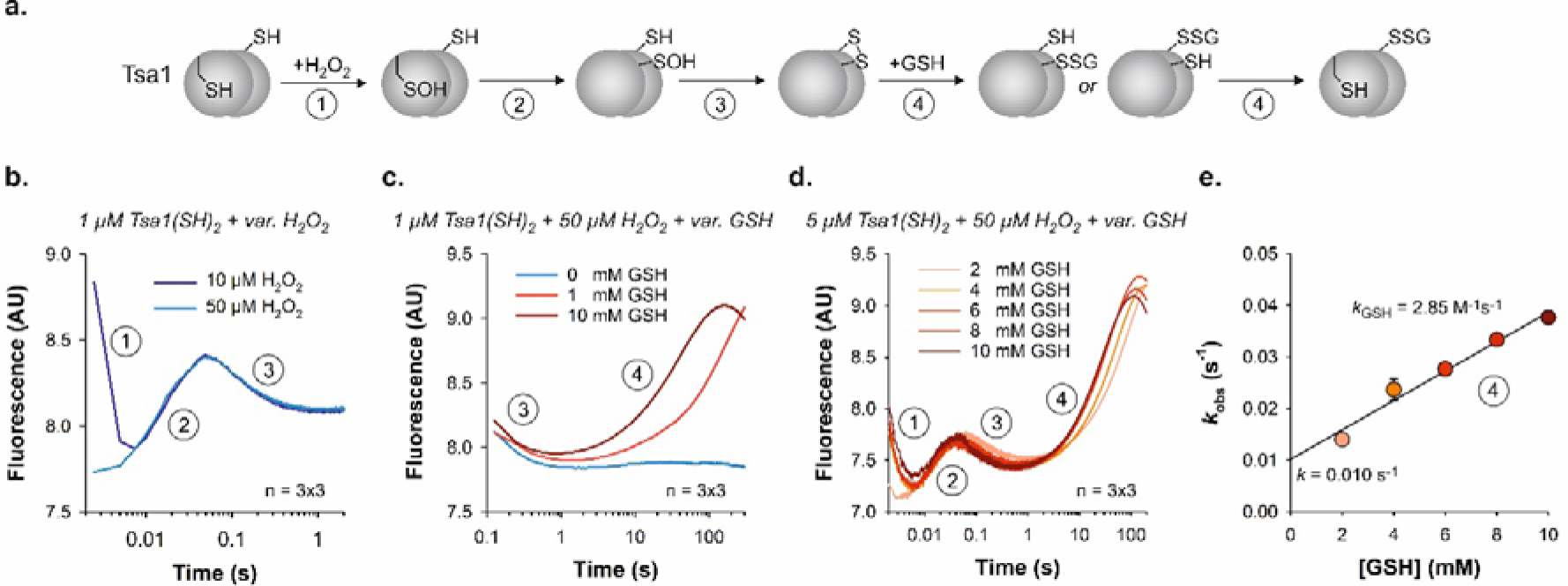
Nonenzymatic reduction of oxidized Tsa1 by GSH. **a.** Reaction scheme for the oxidation of reduced Tsa1 by H_2_O_2_ (step 1), local unfolding (step 2), intramolecular disulfide bond formation (step 3), nonenzymatic glutathionylation and subsequent local refolding of Tsa1 (step 4). **b.** Stopped-flow kinetic measurement of the Tsa1 tryptophan fluorescence for the oxidation of reduced recombinant Tsa1 by H_2_O_2_ at 25°C and pH 7.4. Reaction steps 1-3 were assigned according to the scheme in panel a. The kinetic traces represent averages from technical triplicate measurements. **c,d.** Stopped-flow kinetic measurements as in panel b. with either 1 or 5 µM reduced Tsa1 and variable concentrations of GSH in the first syringe and H_2_O_2_ in the second syringe revealed a GSH-dependent fourth phase. **e**. Secondary plot of the *k*_*obs*_ values (derived from single exponential fits of the fourth phase) against the GSH concentration from panel d. Error bars in panel e represent the standard error for three independent biological replicates.

### Glutaredoxin does not catalyze the glutathionylation of Tsa1 in vitro

Next, we analyzed whether the addition of reduced recombinant Grx2 affects the catalytic cycle and the reaction kinetics (Fig. 6a). Recombinant Grx2, which shares 64% sequence identity with Grx1, was chosen because it accounts for 80% of the physiological glutaredoxin activity in yeast cell extracts and protects yeast against the treatment with H_2_O_2_, FeCl_3_, menadione or paraquat ^61,62^. Considering a potential glutaredoxin-dependent reduction of the Tsa1 disulfide, Tsa1(S_2_), Grx2 might either directly reduce Tsa1 following a dithiol:disulfide exchange mechanism or activate GSH for protein disulfide reduction ^54,63-65^. Although the chosen Grx2 concentration was at least four times higher than the estimated cytosolic concentration of Grx2 ^7^, no additional phases were observed with 10 µM reduced Grx2, indicating that Tsa1(S_2_) does not react directly with Grx2 under the chosen assay conditions (Fig. 6b). In contrast to the reaction with GSH in the absence of Grx2 (Fig. 5), we detected a lag phase between the third and fourth phase when reduced Grx2 and GSH were both added to reduced Tsa1 in the first syringe (Fig. 6b). We interpret the lag phase as a cyclic Grx2-catalyzed deglutathionylation and H_2_O_2_-dependent oxidation of Tsa1 that results in the consumption of H_2_O_2_. These steps are masked during the lag phase because they are much faster than the nonenzymatic glutathionylation of Tsa1. The rate-limiting nonenzymatic glutathionylation and local refolding of Tsa1 only became detectable at the end of the last catalytic cycle. This interpretation is supported by the almost identical GSH-dependent second order rate constant as for the fourth phase without Grx2 (Fig. 5d and Fig.6c). Based on the rate constant and the used concentrations, we estimated that about 20–30 µM H_2_O_2_ were consumed during the 200–330 second lag phase. The difference to the 50 µM H_2_O_2_ used in the assay could be explained by a slightly higher rate constant (e.g., from hyperbolic fits of the secondary plots) or the additional nonenzymatic reduction of H_2_O_2_ by Grx2 and/or GSH ^66^. Stopped-flow experiments with Tsa1(S_2_) in the first syringe and reduced Grx2 and GSH in the second syringe revealed a single-phase increase of fluorescence in accordance with the fourth phase from Fig. 5c,d, again confirming a rate-limiting nonenzymatic reaction with a rate constant *k*_GSH_ of 2.7 M^−1^s^− 1^ despite the presence of Grx2 (Fig. 6d,e). In summary, Grx2 is neither a direct reductant for Tsa1(S_2_) nor activates GSH for Tsa1 reduction, and the rapid Grx2-catalyzed deglutathionylation of Tsa1 is kinetically masked by the rate-limiting nonenzymatic reaction between Tsa1(S_2_) and GSH.

**Figure 6.**
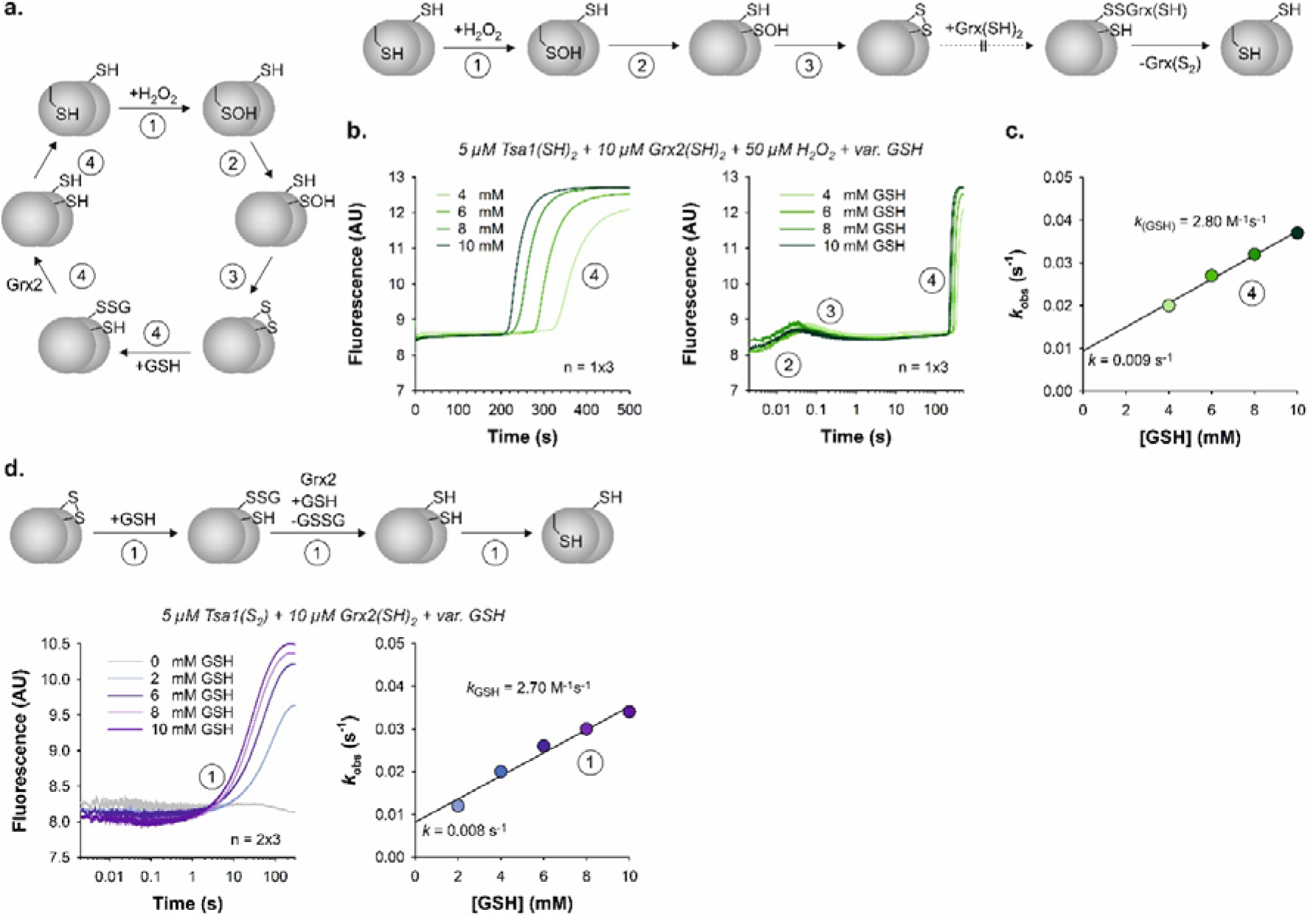
Grx2 reacts with glutathionylated Tsa1 but not with Tsa1(*S*_2_). **a.** Reaction schemes for the reduction of glutathionylated Tsa1 (left side) or of Tsa1(S_2_) (right side) by reduced Grx2. **b.** Stopped-flow kinetic measurement of the Tsa1 tryptophan fluorescence for the oxidation of reduced recombinant Tsa1 by H_2_O_2_ in the presence of Grx2 and variable concentrations of GSH at 25°C and pH 7.4. All reduced compounds were in the first syringe and H_2_O_2_ was in the second syringe. Reaction steps 1-4 were assigned according to the schemes in panel a. **c**. Secondary plot of the *k*_*obs*_ values (derived from single exponential fits of the fourth phase) against the GSH concentration from panel b. **d.** Reaction scheme, stopped-flow kinetic measurement and secondary plot for the reaction between Tsa1(S_2_) in the first syringe and reduced Grx2 and variable concentrations of GSH in the second syringe. The kinetic traces in panels b and d represent averages from technical triplicate measurements. Values in the secondary plots in panels c and d were from single or double biological replicates.

### Nonenzymatic reduction of Tsa1 by GSH would require oxidation of the thioredoxin pool

Recombinant Trx1 was previously shown to reduce Tsa1 in steady-state kinetic assays ^57^). We now determined the rate constants for the Trx1-dependent reduction of Tsa1 using stopped-flow measurements with Tsa1(S_2_) in the first syringe and variable concentrations of reduced Trx1 in the second syringe (Fig. 7a,b). The detected increase in fluorescence could be subdivided into three phases (Fig. 7b). The *k*_*obs*_ values for the strong increase in fluorescence during the first phase depended on the Trx1 concentration (Fig. 7c). A second order rate constant *k*_*Trx1*_ of 2.8×10^6^ M^−1^s^−1^ and a Trx1-independent rate constant of 7.9 s^−1^ were determined from the slope and y-axis intercept of the secondary plot (Fig. 7c). We assign *k*_*Trx1*_ to the reduction of Tsa1(S_2_) yielding a mixed disulfide with Trx1 (whereas the Trx1-independent rate constant might reflect a conformational change or the reverse reaction). The second and third phase appeared to be independent of the Trx1 concentration and remain to be analyzed in more detail. We provisionally assign these two phases to the formation of the Trx1 disulfide, Trx1(S_2_), and the subsequent local refolding of reduced Tsa1 (Fig. 7a,b). Taking into account calculated estimates for the cytosolic concentration of Trx1 between 0.07 and 17 µM ^7^ and GSH around 13 mM ^67^, we then compared the reduction rates for Tsa1(S_2_) (Fig. 7d). At an estimated low Trx1 concentration of 0.07 µM, the reduction of Tsa1 by Trx1 would be about five times faster as for GSH and 20% of Tsa1 might be reduced by GSH. However, this calculation neglects the presence of Trx2, which shares 78% sequence identity with Trx1, has a similar abundance, and also efficiently reacts with Tsa1 ^57^. At an estimated high Trx1 concentration of 17 µM, the reduction of Tsa1 by Trx1 would be about 1.3×10^3^ times faster as for GSH and only 0.08% of Tsa1 might be reduced by GSH (neglecting again the presence of Trx2) (Fig. 7d). To exclude that GSSG is formed by the nonenzymatic reduction of thioredoxins, we also performed control experiments with Trx1(S_2_) and GSH, yielding an estimated second order rate constant < 1 M^−1^s^−1^ (Fig. S7a). Adding either 10 µM Grx2(SH)_2_ or 0.4 U/ml glutathione reductase and NADPH did not accelerate the nonenzymatic GSH-dependent reduction of Trx1(S_2_) (Fig. S7b,c). Thus, Grx2 does not activate GSH for the reduction of Trx1 and a direct reduction of Trx1(S_2_) by GSH is too slow to explain the observed Tsa1-dependent GSSG formation in yeast. In summary, provided that Trx1 and Trx2 are efficiently reduced by thioredoxin reductase, the thioredoxins are most likely the dominant physiological reductants for Tsa1. A nonenzymatic reduction of Tsa1 by GSH, followed by the rapid Grx2-dependent formation of GSSG, requires either a depletion or oxidation of the thioredoxin pool, which probably occurs at high bolus peroxide treatments.

**Figure 7.**
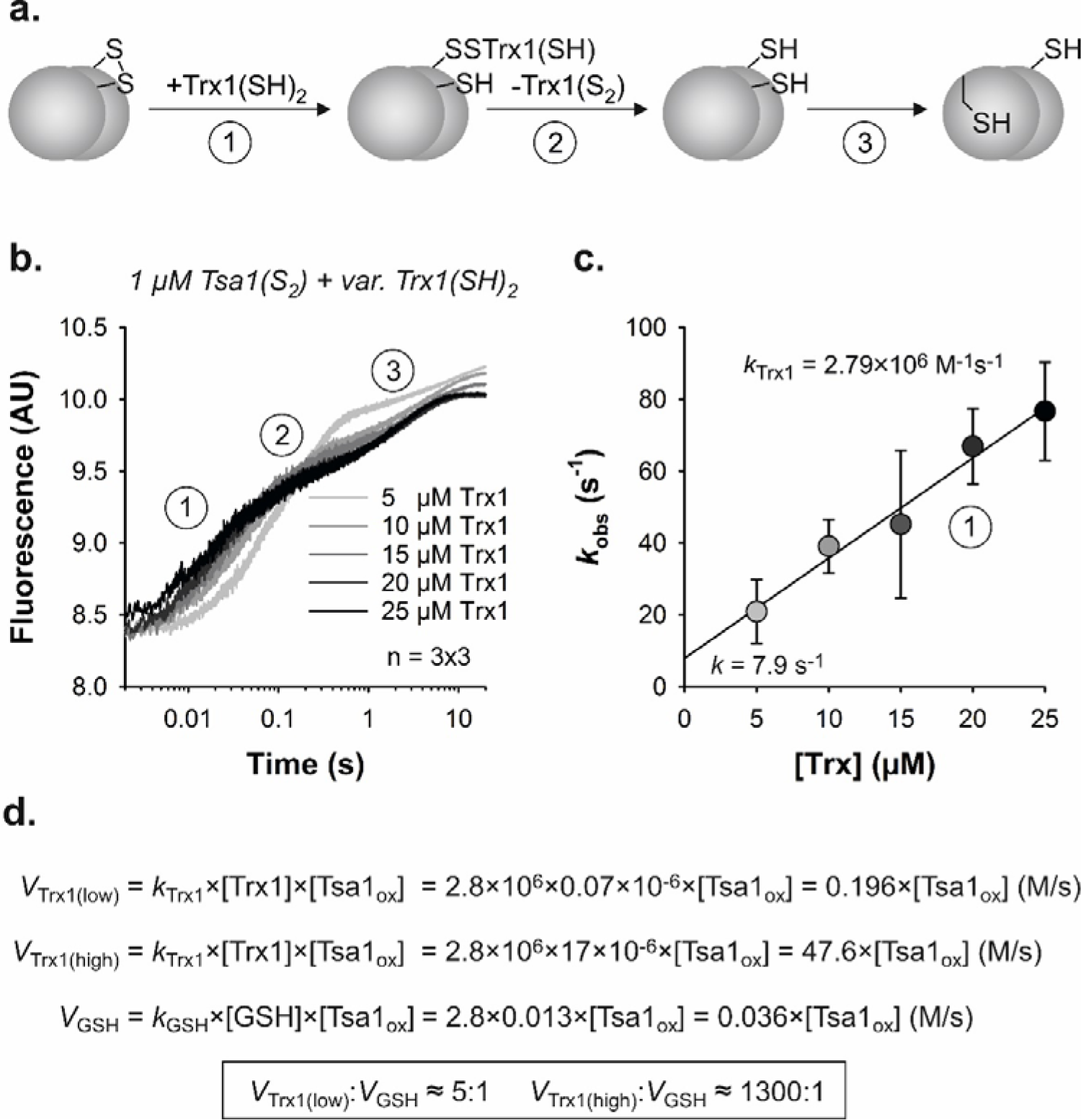
Trx1 rapidly reduces Tsa1(S_2_). **a**. Reaction scheme for the reduction of Tsa1(S_2_) by reduced Trx1. **b**. Stopped-flow kinetic measurement of the Tsa1 tryptophan fluorescence for the reduction of recombinant Tsa1(S_2_) by reduced Trx1 at 25°C and pH 7.4. Reaction steps 1-3 were (provisionally) assigned according to the scheme in panel a. The kinetic traces represent averages from technical triplicate measurements. **c**. Secondary plot of the *k*_*obs*_ values (derived from single exponential fits of the first phase) against the Trx1 concentration from panel b. Error bars in panel c represent the standard error for three independent biological replicates. **d**. Kinetic comparison of the reductions rates for Tsa1(S_2_) by Trx1 or GSH based on estimates for the physiological concentration of the reaction partners.

## Discussion

The efficiency of cellular H_2_O_2_ removal, and the identity of the proteins that facilitate it, has important implications for our understanding of both H_2_O_2_ toxicity and the mechanisms by which H O functions as a signaling molecule ^4,13,68-70^. Our observation that the diffusion of peroxide through the cell is strongly limited by Tsa1 is consistent with previous *in silico* studies, which suggested that peroxiredoxins in general should strongly limit H_2_O_2_ diffusion ^71-73^, studies of budding yeast growth dynamics and the adaptation to H_2_O_2_ challenge ^74^, and by recent observations in fission yeast and mammalian cells, which reported that peroxiredoxin- and thioredoxin reductase-dependent H_2_O_2_ scavengers control intracellular H_2_O_2_ diffusion ^33,75-78^. Taken together, these observations show that steep intracellular H_2_O_2_ gradients do exist and that they are shaped by endogenous thiol peroxidase-dependent scavenging. It will be extremely interesting in the future to test whether intracellular H_2_O_2_ gradients have a role in H_2_O_2_ signaling, for example by helping to ensure specificity. It might be important, that only a specific fraction of a certain protein, in a defined subcellular localization, is oxidized by H_2_O_2_. This could be achieved by co-localization of a peroxiredoxin, target protein and H_2_O_2_ source in specific microdomains, which may be mediated by scaffold proteins ^79,80^, or by a local inactivation of a peroxiredoxin by posttranslational modifications ^81^.

We demonstrate unequivocally that Tsa1 is the dominant scavenger of both H_2_O_2_ and *t*-BuOOH, when applied exogenously at low micromolar concentrations to budding yeast cells. This observation holds true under both fermentative and respiratory conditions. These observations reveal a similarity to the situation previously reported in *Escherichia coli*, where only the alkylhydroperoxide reductase AhpC is relevant for peroxide scavenging at low H_2_O_2_ concentrations ^47,82,83^. *E. coli* harbors other proteins, that display thiol peroxidase activity *in vitro*, but their contribution to *in vivo* H_2_O_2_ detoxification appears negligible ^82^. Nonetheless, the situation is more complex in budding yeast, as it is in most eukaryotes, with budding yeast harboring eight thiol peroxidases, two catalases and a cytochrome *c* peroxidase. Therefore, the bigger question is why cells harbor multiple thiol peroxidases when in many cases only one plays a major role in peroxide removal. The answer to this question remains unclear but may well lie in alternative functions, for example the participation of thiol peroxidases in redox relays, which facilitate the transfer of oxidative equivalents from H_2_O_2_ to specific target proteins to regulate function or activity ^13,34,38,69,70,84,85^. Another explanation could be different substrate preferences or degrees of substrate promiscuity, e.g. for lipid hydroperoxides as electron acceptors or alternative thiols as electron donors. Such properties are indeed reflected by the structural and mechanistic versatility of thiol peroxidases ^3,54,59^. Alternatively, some proteins may be important for peroxide removal in specific subcellular compartments, affording protection in these domains, but having a negligible impact on the overall cellular H_2_O_2_ scavenging capacity.

We show that the two catalases and cytochrome *c* peroxidase in yeast only have a noticeable effect on cellular H_2_O_2_ scavenging capacity when deleted in combination, and even then, only at higher H_2_O_2_ concentrations. This is consistent with the known enzymatic properties of catalases and peroxiredoxins, i.e. catalases cannot be saturated by H_2_O_2_ and do not require a thiol reductant, which allows them to function at very high peroxide concentrations when thioredoxins are oxidized and/or peroxiredoxins are hyperoxidized ^7,33,78^. In terms of H_2_O_2_ toxicity, surprisingly, we still do not fully understand the mechanisms by which H_2_O_2_ leads to cell death, especially in eukaryotes. In prokaryotes, this question has been extensively studied, for example in *E. coli*. The predominant mechanism of H_2_O_2_-induced cell death seems to be indirect damage to DNA via the iron-dependent production of hydroxyl radicals, as well as damage to iron-sulfur clusters ^86,87^. However, in eukaryotes, H_2_O_2_ -dependent oxidative damage does not seem sufficient to fully explain H_2_O_2_ toxicity, particularly when H_2_O_2_ is present at low micromolar to millimolar concentration. Instead, recent studies have pointed to the depletion of pools of reduced thioredoxin and reduced glutathione ^30,31^ as well as modulation of PKA activity as important mediators of H_2_O_2_ -dependent cell death ^35^. Tsa1 was suggested to repress PKA activity in the presence of H_2_O_2_. In a Δ*tsa1* strain, which originally displays a higher sensitivity to H_2_O_2_-dependent cell death than wild-type cells, wild-type-like H_2_O_2_ resilience could be gained by mutations that constitutively repress PKA activity ^35^. Molin and co-workers concluded that whilst ‘*Tsa1 is required for both promoting resistance to H*_2_*O*_2_ *and extending lifespan upon caloric restriction* … *Tsa1 effects both these functions not by scavenging H*_2_*O*_2_*, but by repressing the nutrient signaling Ras-cAMP-PKA pathway at the level of the protein kinase A (PKA) enzyme*’ ^35^. This could thus support the conclusion that the sensitivity of Δ*tsa1* cells to H_2_O_2_ is independent of a loss of H_2_O_2_ scavenging. On the other hand, inactivation of PKA leads to expression of an array of stress resistance enzymes that may increase H_2_O_2_ scavenging capacity, which may not happen as efficiently in the absence of Tsa1. Therefore, whether or not there is a direct link between H_2_O_2_ scavenging capacity and cell death remains to be fully answered.

Finally, we showed that *in vitro* Tsa1 is efficiently reduced by yeast Trx1 but not by Grx2 and/or GSH, suggesting that the intracellular Tsa1-dependent GSSG production requires (i) a depletion or oxidation of the thioredoxin pool and (ii) the presence of a reduced glutaredoxin. The rate constant of 2.8×10^6^ M^−1^s^−1^ for the reduction of Tsa1(S_2_) by yeast Trx1 is very similar to the rate constants for the reduction of the human Prx1-type typical 2 Cys peroxiredoxin PRDX1 by human Trx1 and Trx2 ^60^, the rate constant for the reduction of the Prx1-type typical 2 Cys peroxiredoxin c-TXNPx from *Trypanosoma cruzi* by tryparedoxin ^88^, the rate constant for the reduction of the human Prx5-type atypical 2 Cys peroxiredoxin PRDX5 by human Trx2 ^89^, and the rate constants for the reduction of the sulfenic acid of the promiscuous Prx5-type 1 Cys peroxiredoxin PfAOP from *Plasmodium falciparum* by GSH and other low-molecular-weight thiols ^59,90^. In contrast, the million times smaller rate constant for the reduction of Tsa1(S_2_) by GSH of 2.9 M^−1^s^−1^ indicates a rate-limiting nonenzymatic thiol:disulfide exchange reaction, which is followed by a rapid Grx2-dependent deglutathionylation and GSSG formation. An analogous reaction sequence was previously suggested as one of several mechanisms for the GSH-dependent reduction of human PRDX2 ^55^. In contrast to human PRDX2 ^55^, our measurements do not support a competition between GSH and the resolving cysteinyl residue of the peroxiredoxin for the reduction of the sulfenic acid. The GSH/glutaredoxin system can therefore only serve as a backup system for the thioredoxin system when the yeast thioredoxins are depleted or oxidized. Bolus peroxide treatments of yeast cells could instantaneously oxidize a major fraction of Tsa1. Since the Tsa1 concentration most likely exceeds the concentrations of Trx1 and Trx2 by one or more orders of magnitude, a single catalytic cycle could completely oxidize the thioredoxin pool. Trx1(S_2_) did not react with GSH ^91^. Whether the thioredoxins are efficiently reduced again therefore depends on the availability of NADPH and the concentration and activity of thioredoxin reductase. Our glutathione probe responses indicate that thioredoxin reduction can become limiting during bolus peroxide treatments resulting in the Tsa1- and Grx2-dependent formation of GSSG.

In conclusion, we show that Tsa1 is the dominant scavenger of exogenous H_2_O_2_ and *t*-BuOOH in budding yeast that can become an important source of cytosolic GSSG when the thioredoxin pool becomes oxidized.

## Acknowledgements

B.M., JR and M.D. acknowledge generous support from the Deutsche Forschungsgemeinschaft (DFG) Priority Program SPP1710 (MO 2774/2-1, project number 386433891, and DE 1431/8-2 project number 249669453) and grants MO 2774/7-1, project number 508372800, and DE 1431/19-1 project number 508372800) as well as grants RI2150/5-1 project number 435235019, SPP2453 project number 541742459, RTG2550/1 project number 411422114, and CRC1218 -project number 269925409. LPR acknowledges support from the DFG (SFB/TRR219 project number 322900939). The following yeast strains, BY4742 Δ*ccp1*, BY4742 Δcta1Δctt1, BY4742 Δ*ccp1*Δ*cta1*Δ*ctt1*, BY4742 pRS403, and BY4742 pRS403-*TSA1* were a kind gift from Mikael Molin, Chalmers University of Technology, Gothenburg, Sweden.

## Author contributions

B.M., M.D. and J.R. conceived the study, helped design and supervise the experiments, analyzed and interpreted experimental data and wrote the manuscript. J.Z., L.L., H.L., P.S.A., C.M., T.S., J.O., A.T., T.N.E.O., GC, EP, MW and L.P.R. helped design and perform experiments, analyzed data and helped to write the manuscript.

## Materials and methods

### Growth of yeast cells

All experiments in this study were performed in a *Saccharomyces cerevisiae* BY4742 (MATα *his3*Δ1 *leu2*Δ0 *lys2*Δ0 *ura3*Δ0) or YPH499 (*MAT*a *ura3-52 lys2-801_amber ade2-101_ochre trp1-*Δ*63 his3-* Δ*200 leu2-*Δ*1*) background. Cells were grown in Hartwell’s Complete (HC) medium with 2% glucose or a 2% glycerol / 2% ethanol mixture as carbon sources.

### Construction of yeast strains and plasmid transformation

Most of the yeast gene deletion and gene overexpression strains used here were constructed in previous studies ^36,42^.For the construction of new yeast gene deletion and overexpression strains a standard homologous recombination-based approach was used ^44^. For gene deletion, antibiotic resistance cassettes were amplified by PCR using primers designed to have 50–60 base-pair overhangs homologous to the up- and down-stream regions of the gene to be deleted. Subsequently, the PCR product was transformed into yeast cells using a standard lithium acetate-based protocol. Briefly, cells were harvested and washed with sterile water and resuspended in 200 µl ‘One-Step-Transformation’ buffer consisting of 40% (w/v) polyethylene glycol (PEG), 100 mM dithiothreitol (DTT) and 200 mM lithium acetate together with 10 µl salmon sperm DNA and 10 µl PCR product. Cells were incubated at 45°C for 30 min, transferred to 10 ml of YPD and grown overnight at 30°C. Cells were subsequently streaked onto YPD plates containing the appropriate antibiotic for selection. Successful gene deletions were confirmed by PCR on genomic DNA using primers designed to bind ∼500 bp up- and down-stream of the gene of interest. For genomic DNA extraction, cells were resuspended in 30 µl of 0.2% (w/v) SDS and heated at 95°C for 10 min. Subsequently, cells were briefly vortexed and centrifuged at 18000 *g* for 1 min. 2 µl of supernatant was used as template DNA in the PCR reaction mix.

### Cloning and plasmid construction

All genetically encoded probes used in this study were encoded on p415TEF plasmids. The p415TEF roGFP2-Grx1 ^53^, roGFP2-Tsa2ΔC_R_ ^36^ and Su9-roGFP2-Tsa2ΔC _R_ ^36^ were generated in previous studies. HyPer7 was synthesized with codons optimized for yeast expression (Genscript®) and subcloned into an empty p415TEF plasmid using *XbaI* and *XhoI* restriction sites. The p415TEF HyPer7 plasmid was used as a template for mutation of the H_2_O_2_-reactive cysteine 121 to serine using HyPer7C121S forward primer 5’-CTGAGGGTAACTCTATGAGAGATC-3’ and the HyPer7C121S reverse primer 5’-GATCTCTCATAGAGTTACCCTCAG-3’. Mutation of C121 was performed using a standard site-directed mutagenesis protocol using S7-Fusion Polymerase (*Biozym*). Methylated template DNA was digested by *DpnI* (*NEB*) and 5 µl of the reaction were directly transformed into *E. coli* Top10 cells following plasmid extraction. All plasmids were confirmed by sequencing (Eurofins genomics). Plasmid transformation was performed accordingly. Briefly, cells were harvested, washed and resuspended in 100 µl ‘One-Step-Transformation’ buffer, followed by the addition of 5 µl salmon sperm DNA and 400 ng plasmid DNA. Cells were then incubated with shaking for 30 min at 45°C and subsequently streaked onto HC plates lacking leucine for plasmid selection.

### Intracellular H_2_O_2_ measurements using roGFP2-based redox sensors

For Su9-roGFP2-Tsa2ΔC_R_ measurements, cells were grown in HC media, lacking leucine for plasmid selection with a 2% (v/v) glycerol / 2% (v/v) ethanol mixture as carbon source. For roGFP2-Tsa2ΔC_R_ and roGFP2-Grx1 measurements cells were grown in HC media, lacking leucine for plasmid selection with a 2% glucose (w/v) mixture as carbon source. In all cases, cells were grown at 30°C with shaking until the culture reached a density of *D* ≈ 3.5. Subsequently, cells were resuspended in 100 mM MES/Tris pH 6 buffer to a density of 7.5 *D*_*600*_. Resuspended cells were transferred in 200 µl aliquots to a flat-bottomed 96-well imaging plate (BD Falcon). For every yeast strain measured, one cell aliquot was treated with either 20 mM diamide (*Sigma Aldrich*) and another with 100 mM DTT (*AppChem*) to calibrate full probe oxidation and reduction respectively.

RoGFP2 contains two cysteines located on parallel β-strands adjacent to the chromophore of the GFP. Formation of a disulfide bond between these cysteines induces a change in chromophore protonation and therefore in the excitation spectra. The reduced roGFP2 in its anionic form has a dominant excitation maximum at 488 nm. In contrast, the neutral chromophore in an oxidized roGFP2, exhibits an increased excitation maximum at 400 nm and decreased excitation at 488 nm. Excitation at either 400 or 488 nm leads to fluorescence emission at ∼ 510 nm. The degree of roGFP2 oxidation (OxD) is calculated using the emission intensity at 510 nm following excitation at either 400 nm or 488 nm for the reduced and oxidized controls and the measured sample according to Equation 1.

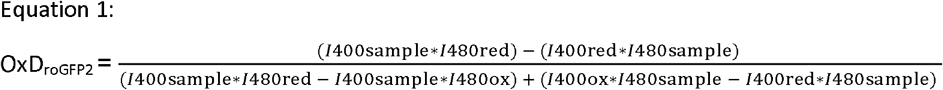

RoGFP2 fluorescence was followed for ∼5 min before the addition of H_2_O_2_ or *t*-BuOOH. The sensor response to the peroxide treatment was followed for ∼30 min using a BMG Labtech CLARIOstar fluorescence plate reader. Cells transformed with an empty p415TEF plasmid were used for fluorescence background subtraction.

### Intracellular H_2_O_2_ measurements using HyPer7

For measurements of HyPer7, cells were grown in HC medium without leucine and 2% (w/v) glucose as carbon source. In all cases, cells were grown at 30°C with shaking until the culture reached a density of *D*_*600*_ ≈ 1.5. Subsequently, cells were resuspended in 100 mM MES/Tris pH 6 buffer to a density of 7.5 *D*_*600*_. Resuspended cells were transferred in 200 µl aliquots to a flat-bottomed 96-well imaging plate (BD Falcon). For every yeast strain measured, one cell aliquot was treated with either 1 mM H_2_O_2_ (*Sigma Aldrich*) and another with 100 mM DTT (*AppChem*) for calibration of the fluorescence plate-reader. HyPer7 is based upon a *circular permutated* yellow fluorescence protein (*cp*YFP) integrated into OxyR domains from *Neisseria meningitidis ^37^*. HyPer7 has two fluorescence excitation maxima depending at 410 nm and 490 nm, which are dominant in the reduced and oxidized states respectively. Excitation at either wavelength results in fluorescence emission at 510 nm.

Fluorescence was measured for ∼5 min before the addition of H_2_O_2_ or *t*-BuOOH. The sensor response to the peroxide treatment was followed for ∼30 min using a BMG Labtech CLARIOstar fluorescence plate reader. Cells transformed with an empty p415TEF plasmid were used for fluorescence background subtraction.

### Purification of recombinant proteins

NADPH was from Gerbu. Diethylenetriaminepentaacetic acid (DTPA), GSH, yeast glutathione reductase (GR), H_2_O_2_, 3-(dimethylcarbamoylimino) -1,1-dimethylurea (diamide) and 1,4-dithiothreitol (DTT) were from Sigma Aldrich. Plasmids pET15b/ScGrx2 and pET15b/Tsa1 encoding MGSSH_6_SSGLVPRGSH-tagged Grx2 and Tsa1 were generated by PCR amplification of the coding sequence from pre-existing plasmids, p415 TEF ScGrx2 ^64^ and p415 TEF roGFP2-Tsa1 ^36^ using the FWD Grx2 primer catgCATATGGTATCCCAGGAAACAGTTGCTCAC and the REV Grx2 primer ctagCTCGAGctattgaaataccggcttcaatatttcag as well as the FWD Tsa1 Primer catgCATATGGTCGCTCAAGTTCAAAAGCAAGCTCCAAC and the REV Tsa1 primer ctagCTCGAGTTATTTGTTGGCAGCTTCGAAGTATTCC. Subsequently PCR products were cloned into an empty pET15b using NdeI and XhoI restriction sites. Please note that the mitochondrial targeting sequence in Grx2 was excluded.

The coding sequence for Trx1 was PCR amplified from yeast genomic DNA using the FWD primer ATGCATCATCACCACCATCACcCCATGGTTACTCAATTCAAAACTGC and reverse primer cgggtaccgagctcCTCGAGTTAAGCATTAGCAGCAATGGCTTGC. Subsequently, the PCR product was cloned into an pTrc99a N-6-His vector using NcoI and XhoI restriction sites. Recombinant Grx2 and Tsa1 were produced in *E. coli* strain SHuffle T7 Express and recombinant Trx1 was produced in *E. coli* strain XL1-Blue. All proteins were purified by Ni-NTA affinity chromatography as described previously for related redoxins ^59,65^. Freshly Ni-NTA purified recombinant Tsa1, Grx2 and Trx1 were reduced with 5 mM DTT on ice for 30 min. Excess DTT and imidazole were removed on a PD-10 desalting column (Merck), and the reduced proteins were eluted with 3.5 mL ice-cold assay buffer containing 100 mM Na_x_H_y_PO_4_, 0.1 mM DTPA, pH 7.4 (at 25°C). Reduced Tsa1 or Trx1 was oxidized with 5 mM diamide on ice for 60 min yielding Tsa1(S_2_) or Trx1(S_2_). Excess diamide was removed on a Ni-NTA column before the removal of imidazole on a PD-10 column. All enzymes and reactants were diluted/dissolved in ice-cold assay buffer. The purity of all proteins was confirmed by analytical SDS-PAGE. Protein concentrations of the eluates were determined spectrophotometrically using the following extinction coefficients ε_280 nm_ as calculated at http://web.expasy.org/protparam: 23.95 mM^-1^cm^-1^ for reduced Tsa1, 24.08 mM^-1^cm^-1^ for oxidized Tsa1, 4.47 mM^-1^cm^-1^ for reduced Grx2 and 9.97 mM^-1^cm^-1^ for reduced Trx1. Average yields were 0.4 µmol of recombinant protein per 1 L of *E. coli* culture.

### Stopped-flow kinetic measurements

Recombinant Tsa1, Grx2 and Trx1 were freshly purified, reduced/oxidized and desalted in assay buffer as described above. Stopped-flow measurements were performed at 25°C in a thermostatted SX-20 spectrofluorometer (Applied Photophysics). The change of tryptophan fluorescence was measured for up to 500 s after mixing (total emission at an excitation wavelength of 295 nm with a slit width of 2 mm). The nonenzymatic reduction of *in situ*-oxidized Tsa1 by GSH was investigated by mixing 2 or 10 µM reduced Tsa1 and variable concentrations of GSH in syringe 1 with 20 or 100 µM H_2_O_2_ in syringe 2. For the reduction of *in situ*-oxidized Tsa1 by GSH and reduced Grx2, 10 µM reduced Tsa1 with 20 µM reduced Grx2 and variable concentrations of GSH in syringe 1 were mixed with 100 µM H_2_O_2_ in syringe 2. Alternatively, 10 µM Tsa1(S_2_) in syringe 1 was mixed with 20 µM reduced Grx2 and variable concentrations of GSH in syringe 2. The reduction of Tsa1 by Trx1 was measured by mixing 2 µM Tsa1(S_2_) in syringe 1 with variable concentrations of reduced Trx1 in syringe 2. The reduction of Trx1 was analyzed by mixing 10 µM Trx1(S_2_) in syringe 1 with 20 mM GSH with or without 20 µM reduced Grx2 or 0.8 U/mL GR and 500 µM NADPH in syringe 2. The kinetic traces of three consecutive measurements were averaged and fitted using the Pro-Data SX software (Applied Photophysics). Rate constants *k*_*obs*_ from single exponential fits were plotted against the substrate concentration in SigmaPlot 13.0 to obtain second order rate constants from the slopes of the linear fits.

