## Supplementary Information for "Tsa1 is the dominant peroxide scavenger and a source of H_2_O_2_-dependent GSSG production in yeast"

Prof. Dr. Bruce Morgan

Prof. Dr. Jan Riemer

Prof. Dr. Marcel Deponte

Keywords: roGFP2, HyPer7, peroxiredoxin, catalase, thiol peroxidase, heme peroxidase, H_2_O_2_ scavenging.

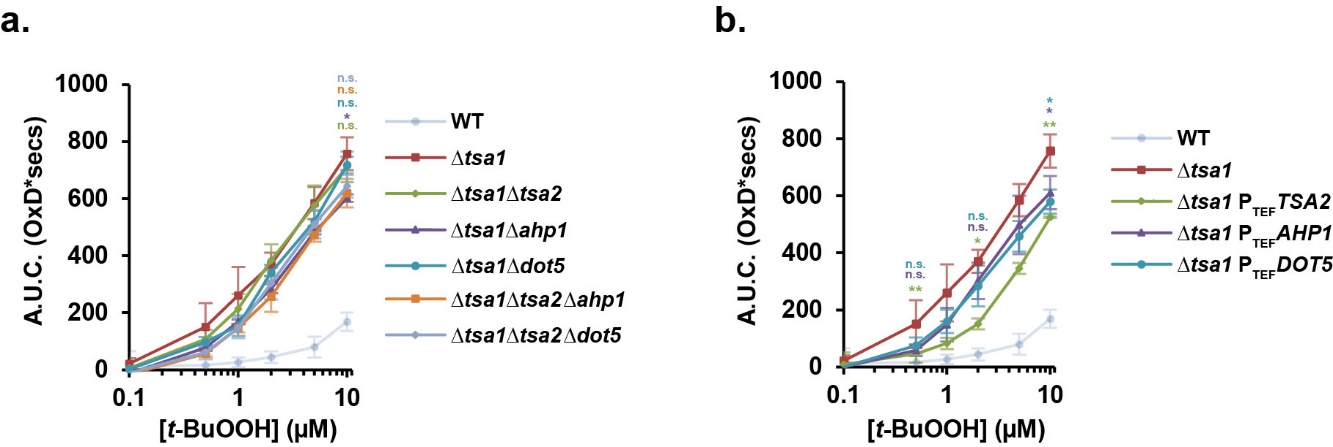

**Supplementary Figure 1. Tsa1 is the dominant scavenger of *t*-BuOOH.**

The area under the curve (A.U.C) of Su9-roGFP2-Tsa2ΔC_R_ probe response to the indicated concentrations of *t*-BuOOH in wild-type cells and ∆*tsa1* cells, together with cells **a.** deleted for, and **b.** overexpressing *TSA2*, *AHP1* and *DOT5*. The WT and Δ*tsa1* data are replicated in both panels. Error bars in all panels represent the standard error for at least three independent repeats. P-values were determined by one-way ANOVA analysis followed by a Tukey’s test.

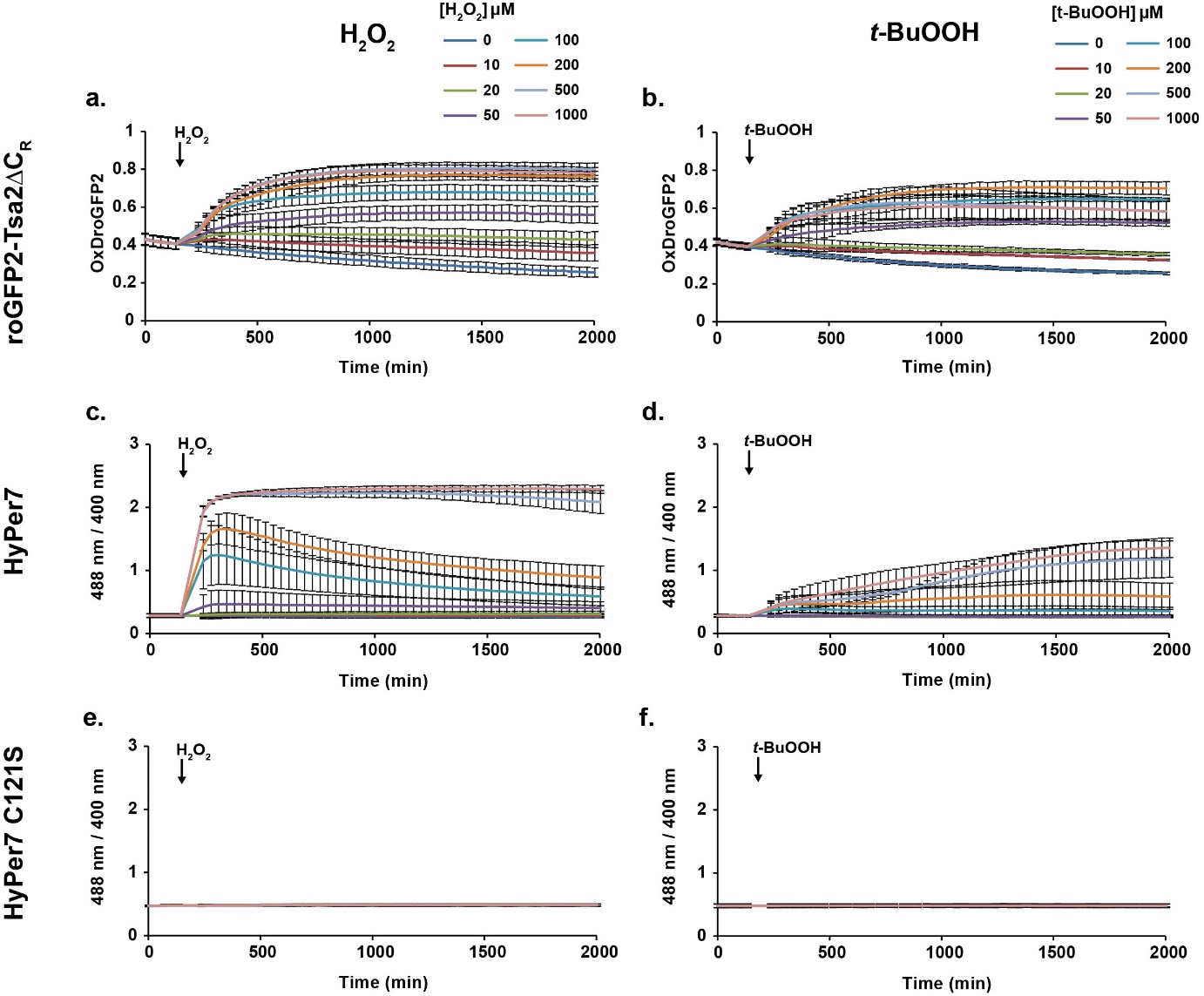

**Supplementary Figure 2. HyPer7 responds to H_2_O_2_ but is poorly responsive to *t*-BuOOH.**

BY4742 wild-type cells were transformed with p415TEF plasmids containing genes for expression of **a.,b.** cytosolic roGFP2-Tsa2∆C_R_, **c.,d.** HyPer7 or **e.,f.** HyPer7C121S. Cells were resuspended in 100 mM MES/Tris pH 6, transferred to a flat-bottomed 96-well plate and subsequently treated at the indicated timepoint with either **a.,c.,e.** H_2_O_2_ or **b.,d.,f.** *t*-BuOOH at concentrations ranging from 10–1000 µM. Error bars in all panels represent the standard error for at least three independent repeats.

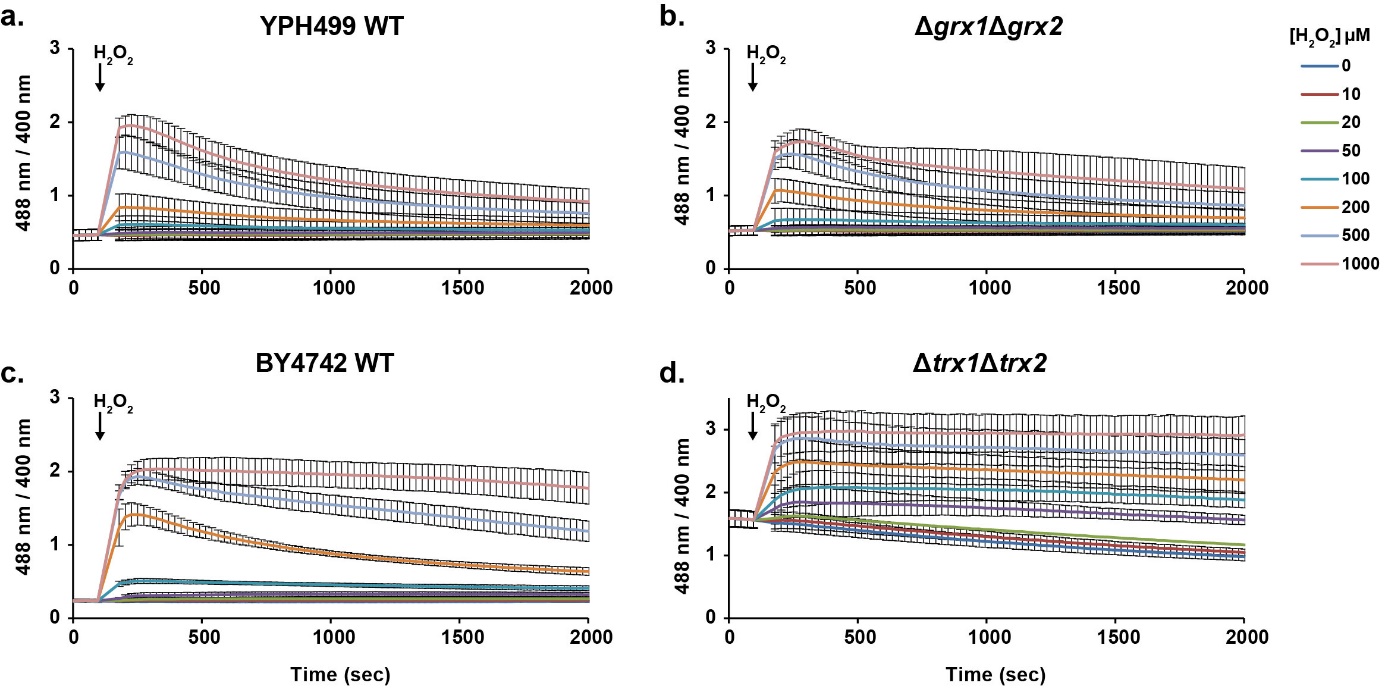

**Supplementary Figure 3. Endogenous reduction of HyPer7 depends upon thioredoxins.**

Response of cytosolic HyPer7 in **a.** YPH499 wild-type, **b.** YPH499 ∆*grx1*∆*grx2*, **c.** BY4742 wild-type and **d.** BY4742 ∆*trx1*∆*trx2* cells treated with exogenously added H_2_O_2_ at concentrations ranging from 10–1000 µM. Error bars in all panels represent the standard error for at least three independent repeats.

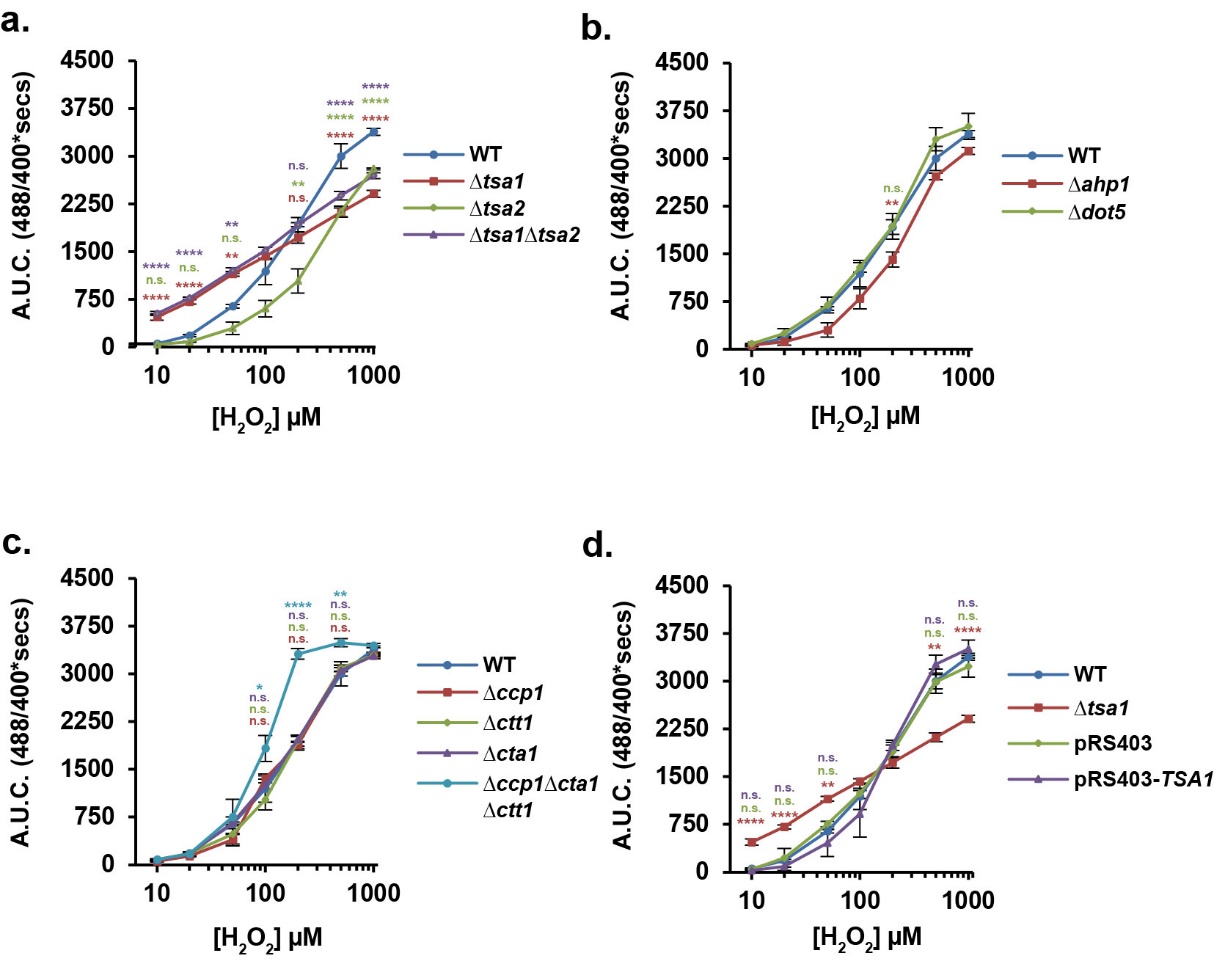

**Supplementary Figure 4. Cytosolic HyPer7 supports a dominant importance of Tsa1 for cytosolic H_2_O_2_ scavenging in cells grown in non-fermentable medium**

**a–d.** HyPer7 responses were measured in the indicated yeast deletion strains in response to exogenous H_2_O_2_ applied at the indicated concentrations. The integrated A.U.C. was calculated for the first 2000 secs of the probe response. The data for wild-type (WT) cells are replicated in each panel. The data for Δ*tsa1* cells is replicated in panels **a**. and **d**. Error bars in both panels represent the standard error of the mean for at least three independent repeats. P values were determined by a one-way ANOVA analysis followed by a Tukey’s (HSD) test.

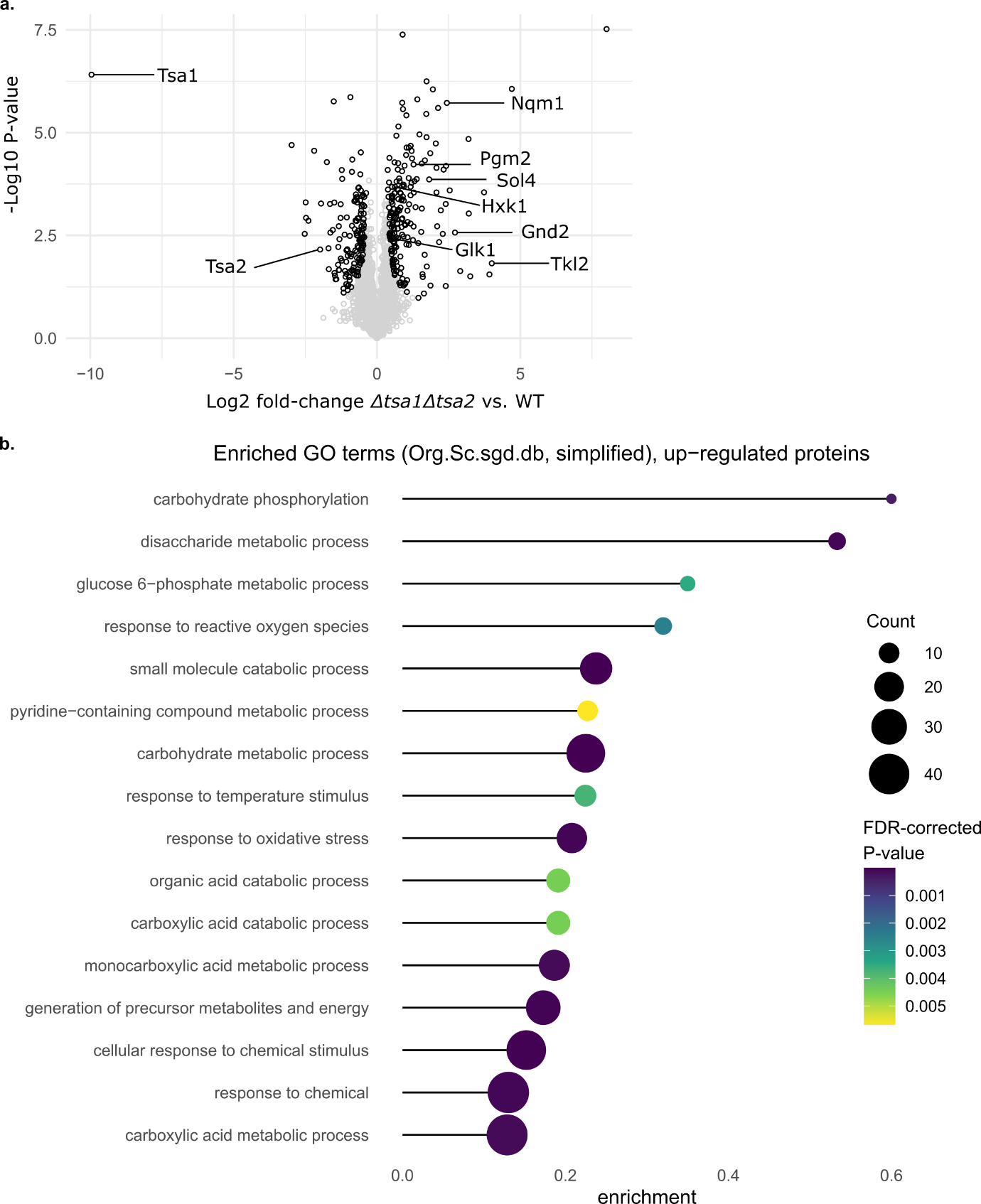

**Supplementary Figure 5. Tsa1 deletion induces compensatory upregulation of pentose phosphate pathway enzymes**

**a.** Results of label-free quantitative proteomics comparing protein abundances in BY4742 ∆*tsa1*∆*tsa2* against BY4742 wild-type (WT). Significantly up- or down-regulated proteins (permutation based-threshold as determined using Perseus [https://maxquant.net/perseus/] with parameters s = 0.2, q = 0.05) are shown in black, others in grey. Proteins belonging to the GO term “glucose-6-phosphate metabolic process” and Tsa1, Tsa2 are highlighted. **b.** GO term overrepresentation analysis was performed for significantly up-regulated proteins as shown in panel a using the clusterProfiler R package (Wu et al., 2021). Redundant GO terms were removed using the ‘simplify’ function (similarity cutoff: 0.7).

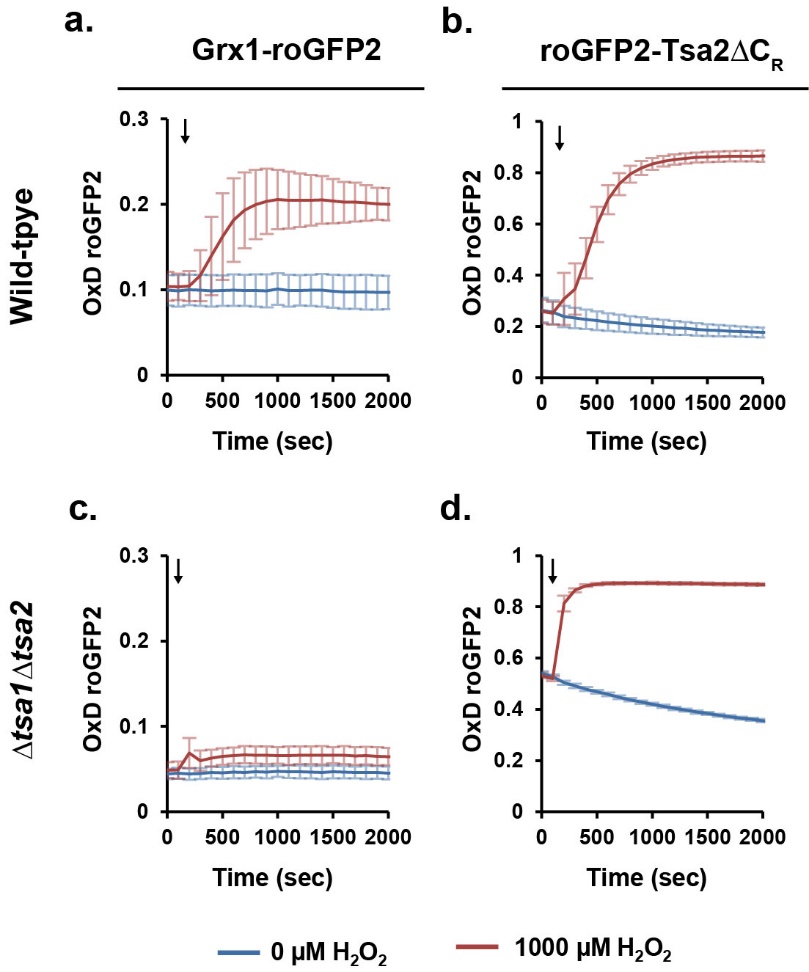

**Supplementary Figure 6. Tsa1 is a major source of cytosolic H_2_O_2_-dependent GSSG.**

**a,c.** Grx1-roGFP2 and **b,d** roGFP2-Tsa2ΔC_R_ probe responses to 0 µM or 1000 µM H_2_O_2_  were measured in **a,b** wild-type and **c,d.** Δ*tsa1*Δ*tsa*2 yeast cells. Error bars represent the standard error of the mean for at least three independent repeats.

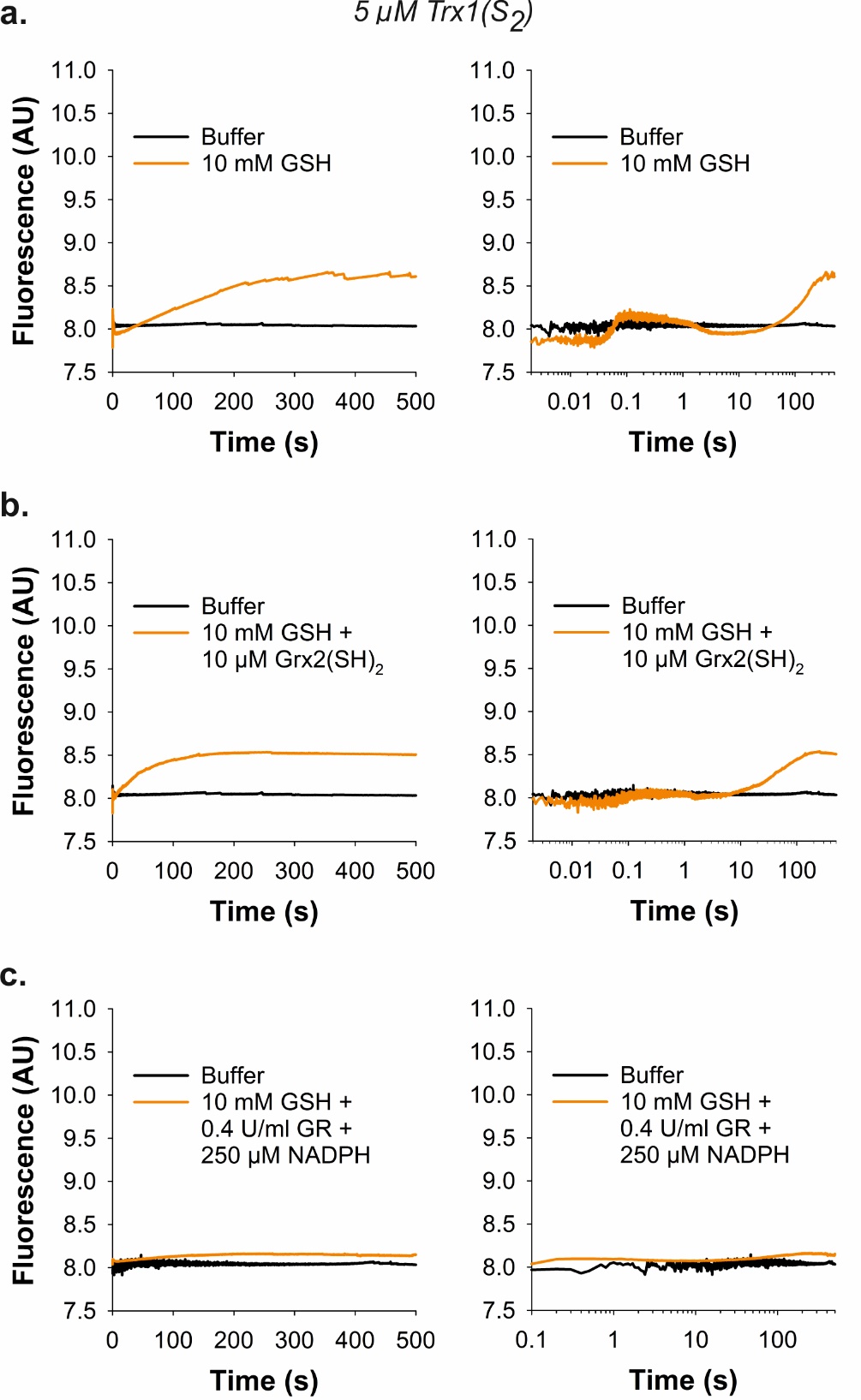

**Supplementary Figure 7. Oxidized Trx1 neither reacts with GSH nor Grx.**

Stopped-flow kinetic measurement of the tryptophan fluorescence for the reduction of recombinant Trx1(S_2_) by **a.** GSH, **b.** GSH and reduced Grx2, and **c.** GSH and GR in the presence of NADPH at 25°C and pH 7.4.

**Supplementary Table 1. Yeast strains used in this study**

Most yeast deletion and overexpression strains used in this work were generated in previous studies unless otherwise indicated. Strains were transformed in this study, with plasmids for probe expression as detailed below.

| Genotype | Figure |
| --- | --- |
| BY4742 *MAT*α *his3*∆*1 leu2*∆*1 lys2*∆*0 ura3*∆*0* + p415TEF empty | 1c-j, 2, 3, 4, 5, S1, S3c,d, S4, S6 |
| BY4742 *MAT*α *his3*∆*1 leu2*∆*1 lys2*∆*0 ura3*∆*0* + p415TEF Su9 roGFP2-Tsa2ΔC_R_ | 1c-j, 2, S1 |
| BY4742 Δ*tsa1::kanMX4* + p415TEF Su9 roGFP2-Tsa2ΔC_R_ | 1c,f,i,j, 2, S1 |
| BY4742 Δ*tsa2::kanMX4* + p415TEF Su9 roGFP2-Tsa2ΔC_R_ | 1c,f |
| BY4742 Δ*tsa1::natNT2* Δ*tsa2::kanMX4* + p415TEF Su9 roGFP2-Tsa2ΔC_R_ | 1c,f, 2a,b, S1a |
| BY4742 Δ*ahp1::kanMX4* + p415TEF Su9 roGFP2-Tsa2ΔC_R_ | 1d,g |
| BY4742 Δ*dot5::kanMX4* + p415TEF Su9 roGFP2-Tsa2ΔC_R_ | 1d,g |
| BY4742 Δ*gpx1::kanMX41* + p415TEF Su9 roGFP2-Tsa2ΔC_R_ | 1d,g |
| BY4742 Δ*gpx2::kanMX4* + p415TEF Su9 roGFP2-Tsa2ΔC_R_ | 1d,g |
| BY4742 Δ*gpx3::kanMX43* + p415TEF Su9 roGFP2-Tsa2ΔC_R_ | 1d,g |
| BY4742 Δ*ctt1::kanMX4* + p415TEF Su9 roGFP2-Tsa2ΔC_R_ | 1e,h |
| BY4742 Δ*cta1::kanMX4* + p415TEF Su9 roGFP2-Tsa2ΔC_R_ | 1e,h |
| BY4742 *his3*∆*1::pRS403* + p415TEF Su9 roGFP2-Tsa2ΔC_R_ | 1i,j |
| BY4742 *his3*∆*1::pRS403-TSA1* + p415TEF Su9 roGFP2-Tsa2ΔC_R_ | 1i,j |
| BY4742 Δ*tsa1::kanMX4* Δ*ahp1::hphNT1* + p415TEF Su9 roGFP2-Tsa2ΔC_R_ | 2a,b, S1a |
| BY4742 Δ*tsa1::kanMX4* Δ*dot5::hphNT1* + p415TEF Su9 roGFP2-Tsa2ΔC_R_ | 2a,b, S1a |
| BY4742 Δ*tsa1::natNT2* Δ*tsa2::kanMX4* Δ*ahp1::hphNT1* + p415TEF Su9 roGFP2-Tsa2ΔC_R_ | 2a,b, S1a |
| BY4742 Δ*tsa1::natNT2* Δ*tsa2::kanMX4* Δ*dot5::hphNT1* + p415TEF Su9 roGFP2-Tsa2ΔC_R_ | 2a,b, S1a |
| BY4742 Δ*tsa1::hphNT1 kanMX4-P_TEF_TSA2* + p415TEF Su9 roGFP2-Tsa2ΔC_R_ | 2c,d, S1b |
| BY4742 Δ*tsa1::hphNT1 kanMX4-P_TEF_AHP1* + p415TEF Su9 roGFP2-Tsa2ΔC_R_ | 2c,d, S1b |
| BY4742 Δ*tsa1::hphNT1 kanMX4-P_TEF_DOT5* + p415TEF Su9 roGFP2-Tsa2ΔC_R_ | 2c,d, S1b |
| BY4742 Δ*ccp1::kanMX4* + p415TEF Su9 roGFP2-Tsa2ΔC_R_ | 2e,f |
| BY4742 Δ*ctt1::natMX4* Δ*cta1::hphMX4* + p415TEF Su9 roGFP2-Tsa2ΔC_R_ | 2e,f |
| BY4742 Δ*ccp1::natMX4* Δ*ctt1::hphMX4* Δ*cta1::kanMX4* + p415TEF Su9 roGFP2-Tsa2ΔC_R_ | 2e,f |
| BY4742 *MAT*α *his3*∆*1 leu2*∆*1 lys2*∆*0 ura3*∆*0* + p415TEF HyPer7 | 3, S2c,d, S3, S4 |
| BY4742 Δ*tsa1::kanMX4* + p415TEF HyPer7 | 3a,d, S4a,d |
| BY4742 Δ*tsa2::kanMX4* + p415TEF HyPer7 | 3a, S4a |
| BY4742 Δ*tsa1::natNT2* Δ*tsa2::kanMX4* + p415TEF HyPer7 | 3a, S4a |
| BY4742 Δ*ahp1::kanMX4* + p415TEF HyPer7 | 3b, S4b |
| BY4742 Δ*dot5::kanMX4* + p415TEF HyPer7 | 3b, S4b |
| BY4742 Δ*gpx1::kanMX41* + p415TEF HyPer7 | 3b |
| BY4742 Δ*gpx2::kanMX4* + p415TEF HyPer7 | 3b |
| BY4742 Δ*gpx3::kanMX43* + p415TEF HyPer7 | 3b |
| BY4742 Δ*ccp1::kanMX4* + p415TEF HyPer7 | 3c, S4c |
| BY4742 Δ*ctt1::kanMX4* + p415TEF HyPer7 | 3c, S4c |
| BY4742 Δ*cta1::kanMX4* + p415TEF HyPer7 | 3c, S4c |
| BY4742 Δ*ccp1::natMX4* Δ*ctt1::hphMX4* Δ*cta1::kanMX4* + p415TEF HyPer7 | 3c, S4c |
| BY4742 *his3*∆*1::pRS403* + p415TEF HyPer7 | 3d, S4d |
| BY4742 *his3*∆*1::pRS403-TSA1* + p415TEF HyPer7 | 3d, S4d |
| BY4742 *MAT*α *his3*∆*1 leu2*∆*1 lys2*∆*0 ura3*∆*0* + p415TEF roGFP2-Tsa2ΔC_R_ | S2a,b, S6b |
| BY4742 *MAT*α *his3*∆*1 leu2*∆*1 lys2*∆*0 ura3*∆*0* + p415TEF HyPer7C121S | S2e,f |
| YPH499 *MAT*α *ura3-52 lys2-801_amber ade2-101_ochre trp1-Δ63 his3-Δ200 leu2-Δ1*  + p415TEF empty | S3a,b |
| YPH499 *MAT*α *ura3-52 lys2-801_amber ade2-101_ochre trp1-Δ63 his3-Δ200 leu2-Δ1*  + p415TEF HyPer7 | S3a |
| YPH499 Δ*grx1::kanMX4* Δ*grx2::URA3* + p415TEF HyPer7 | S3b |
| BY4742 Δ*trx1::kanMX4* Δ*trx2::HIS3* + p415TEF HyPer7 | S3d |
| BY4742 Δ*glr1::kanMX4*  + p415TEF empty | 4 |
| BY4742 Δ*glr1::kanMX4*  + p415TEF roGFP2-Grx1 | 4 |
| BY4742 Δ*glr1::kanMX4* Δ*tsa1::hphNT1*  + p415TEF roGFP2-Grx1 | 4 |
| BY4742 Δ*glr1::kanMX4* Δ*tsa2::hphNT1*  + p415TEF roGFP2-Grx1 | 4 |
| BY4742 Δ*glr1::kanMX4* Δ*tsa1::hphNT1* Δ*tsa2::natNT2* + p415TEF roGFP2-Grx1 | 4 |
| BY4742 *MAT*α *his3*∆*1 leu2*∆*1 lys2*∆*0 ura3*∆*0* + p415TEF Grx1-roGFP2 | S6a |
| BY4742 Δ*tsa1::natNT2* Δ*tsa2::kanMX4* + p415TEF Grx1-roGFP2 | S6c |
| BY4742 Δ*tsa1::natNT2* Δ*tsa2::kanMX4* + p415TEF roGFP2-Tsa2 ΔC_R_ | S6d |
